# Cell-to-cell and genome-to-genome variability of Adenovirus transcription tuned by the cell cycle

**DOI:** 10.1101/2020.02.27.967471

**Authors:** Maarit Suomalainen, Vibhu Prasad, Abhilash Kannan, Urs F. Greber

## Abstract

In clonal cultures, not all cells are equally susceptible to virus infection. Underlying mechanisms of infection variability are poorly understood. Here, we developed image-based single cell measurements to scrutinize the heterogeneity of adenovirus (AdV) infection. AdV delivers, transcribes and replicates a linear double-stranded DNA genome in the nucleus. We measured the abundance of viral transcripts by single-molecule RNA fluorescence in situ hybridization (FISH), and the incoming ethynyl-deoxy-cytidine (EdC)-tagged viral genome by copper(I)-catalyzed azide-alkyne cycloaddition (click) reaction. The early transcripts increased from 2-12 hours, the late ones from 12-23 hours post infection (pi), indicating distinct accumulation kinetics. Surprisingly, the expression of the immediate early transactivator gene E1A only moderately correlated with the number of viral genomes in the cell nucleus, although the incoming viral DNA remained largely intact until 7 hours pi. Genome-to-genome heterogeneity was found at the level of viral transcription, as indicated by colocalization with the large intron containing early region E4 transcripts, uncorrelated to the multiplicity of incoming genomes in the nucleus. In accordance, individual genomes exhibited heterogeneous replication activity, as shown by single-strand DNA-FISH and immunocytochemistry. These results indicate that the variability in viral gene expression and replication are not due to defective genomes but due to host cell heterogeneity. By analyzing the cell cycle state, we found that G1 cells exhibited the highest E1A expression, and significantly increased the correlation between E1A expression and viral genome copy numbers. This combined image-based single molecule procedure is ideally suited to explore the cell-to-cell variability in viral infection, including transcriptional activators and repressors, RNA splicing mechanisms, and the impact of the 3-dimensional nuclear topology on gene regulation.

**Author Summary:** Adenoviruses (AdV) are ubiquitous pathogens in vertebrates. They persist in infected people, and cause unpredictable outbreaks, morbidity and mortality across the globe. Here we report that the common human AdV type C5 (AdV-C5) gives rise to considerable infection variability at the level of single cells in culture, and that a major underlying reason is the cell-to-cell heterogeneity. By combining sensitive single molecule in situ technology for detecting the incoming viral DNA and newly synthesized viral transcripts we show that viral gene expression is heterogeneous between infected human cells, as well as individual genomes. We report a moderate correlation between the number of viral genomes in the nucleus and immediate early E1A transcripts. This correlation is increased in the G1 phase of the cell cycle, where the E1A transcripts were found to be more abundant than in any other cell cycle phase. Our results demonstrate the importance of cell-to-cell variability measurements for understanding transcription and replication in viral infections.

## Introduction

Virus infections have variable outcomes, they can be lytic, chronic, persistent, latent or abortive. For example, lytic infections kill the host cell and release large numbers of progeny particles, whereas chronic infections continuously release infectious particles, for example hepatitis B virus infections of liver hepatocytes [1]. Latent infections do not produce infectious particles, as shown with herpes viruses [2], and persistent infections give raise to low levels of progeny. Infection outcome is important as it determines the severity of disease, yet, the underlying mechanisms are largely complex. They not only depend on cell autonomous factors, such as the type and the innate immune status of the infected cell, but also on the environment, pro- and anti-viral agents, the immune status of the organism or the nature of the inoculum [3, 4]. A switch between lytic and persistent infection has for example been documented for human adenovirus (AdV) infection of human diploid fibroblasts in presence of type I or II interferon (IFN) due to transcriptional silencing of the E1A enhancer/promoter [5]. AdV are non-enveloped, double-stranded DNA viruses that cause mild respiratory infections in immuno-competent hosts, and establish persistent infections, which can develop into life-threatening infections if the host becomes immuno-compromised [reviewed in 6]. In AdV persistence or latency in human T-lymphocytes and tonsils, the viral genome has no detectable transcriptional activity of E1, E2 and late genes, nor virus production [7–9].

In principle, the cell autonomous aspects of the variable infection outcomes can be addressed by studying cell-to-cell variable parameters with purified virus inocula in defined cell types. For example, the cell-to-cell variable viral transcript counts and progeny yields have been observed with influenza, lymphocytic choriomeningitis, arenavirus, foot-and-mouth disease virus, Herpes simplex virus 1, murine gamma-herpesvirus 68 and Dengue and Zika virus [10–17]. Studies with Influenza A virus revealed variability in translation and assembly of progeny particles [18, 19].

Studies with AdV show further cell-to-cell heterogeneity, namely in virion binding to the cells [20], endosomal and cytoplasmic trafficking [21, 22], and virion uncoating [23–28]. Single-cell, single-particle analyses show that different steps of entry occur with different efficiencies in individual infected cells and thus the number of vDNAs delivered into the nucleus varies between cells [21, 25, 27]. AdV enters cells by receptor-mediated endocytosis, and receives chemical and mechanical cues which initiate the stepwise uncoating process of the viral DNA (vDNA) genome, and delivers vDNA into the nucleus (reviewed in [29–32]). The vDNA is imported into the nucleus in association with protein VII and the protein VII-complexed vDNA is the template for early virus transcription [25, 33–38], although partial replacement of protein VII by histones cannot be excluded [39]. Two other vDNA-associated proteins are protein V and protein X/µ, but in contrast to protein VII, protein V dissociates from the incoming genome before nuclear import, and the fate of protein X/µ is unknown [40].

So far, AdV gene expression has been studied by classical cell population-level assays, such as Northern blots, quantitative reverse transcription polymerase chain reaction (qRT-PCR), microarrays or bulk cell RNA-sequencing [41, 42]. Following nuclear import of the AdV-C vDNA, an enhancer sequence on the left-end of the viral genome activates the host RNA polymerase II-mediated transcription from the E1A transcription unit promoter [43]. Alternative splicing produces initially two different mRNAs, the 13S and 12S E1A mRNAs [44, 45], which yield 289- and 243-residue multifunctional proteins interacting with numerous cellular partners and dramatically changing the host cell [reviewed in 46, 47]. For example, E1A proteins remodel host gene expression, induce the cell to enter the S-phase of cell cycle, suppress host innate immune responses and contribute to reprogramming of cell metabolism. In addition, the 289-residue E1A protein stimulates transcription from its own promoter and activates transcription from the other viral early transcription units, the E1B, E2, E3 and E4, each of which yield multiple mRNAs due to alternative splicing [48–52]. Transcripts from the viral late transcription unit, amongst them mRNAs for the viral structural proteins, become abundant after vDNA replication [52–55]. Our single-cell and single vDNA and RNA resolution assays further address the cell-to-cell variability of AdV infection. They reveal a highly asynchronous and heterogeneous accumulation of viral early transcripts at the onset of viral protein translation, not due to defective viral genomes but partly due to cell cycle effects.

## Results

### Visualization of AdV-C5 transcripts in single cells

We used fluorescent in situ hybridization (FISH) with probes targeting E1A, E1B-55K and protein VI transcripts followed by branched DNA (bDNA) signal amplification to visualize the appearance and abundance of viral transcripts in AdV-C5-infected A549 lung carcinoma cells. This enables analysis of mRNAs at single-cell and single-molecule resolution [56, 57]. AdV-C5 was added to cells at 37°C for 60 min at multiplicity of infection (moi) of ∼54400 virus particles per cell. After removal of unbound virus, cells were incubated at 37°C for 4 h or 11 h before fixation, staining and imaging by confocal microscopy. This procedure ensures that infection is initiated in a relatively short time window, 60 min. It enables estimation of virus particles bound to cells in a parallel sample. The virus particles bound to cells were visualized by mouse anti-hexon 9C12 and anti-mouse Alexa Fluor488-conjugated antibodies and counted from maximum projections of confocal image stacks, amounting between 6 and 173 particles per cell (median 75; Fig. 1A).

**Figure 1:**
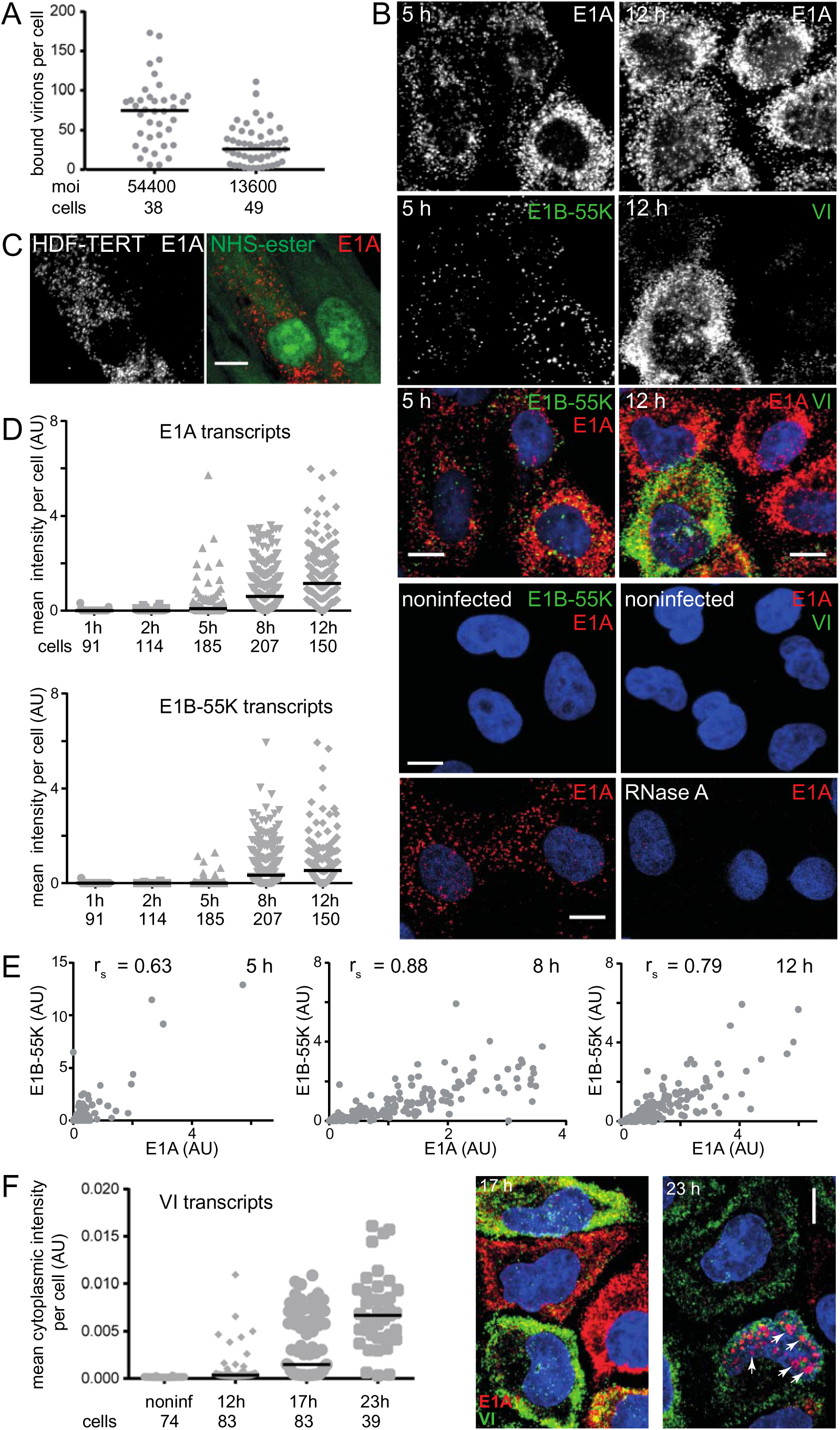
Visualization of AdV-C5 E1A, E1B-55K and protein VI transcripts in infected cells by bDNA-FISH technique. A) Correlation between input virus amounts and number of virus particles bound to cells. Virus was added to A549 cells at 37°C for 60 min (moi ∼ 54400 or 13600 virus particles per cell). Cells were fixed after removal of unbound virus, stained with mouse 9C12 anti-hexon and Alexa Fluor488-conjugated anti-mouse antibodies and imaged by confocal microscopy. The scatterplot shows number of virus particles per cell, one dot representing one cell. Horizontal bars represent median values and the number of cells analyzed per sample is indicated. B) Infected cells accumulate high numbers of viral transcripts. A549 cells were incubated with virus at moi ∼ 54400 virus particles per cell as described in (A) and fixed after 4 h (left-hand panel) or 11 h (right-hand panel) post removal of unbound virus. Fixed infected and noninfected cells were stained with probes against E1A and E1B-55K mRNAs (left-hand panel) or E1A and protein VI mRNAs (right-hand panel). RNase A treatment prior to staining removes transcript signals, as shown for E1A. C) High number of E1A transcripts can be detected in infected HDF-TERT cells. AdV-C5 was added to cells at 37°C for 12 h (moi ∼ 37500 virus particles per cell). After removal of unbound virus, incubation was continued at 37°C for additional 10 h before cells were fixed and stained for E1A mRNAs. Alexa Fluor647 NHS Ester was used for staining of cell area (pseudo-colored green in the figure). D) Timecourse analysis of E1A and E1B-55K transcript accumulation. Virus was added to A549 cells at 37°C for 60 min (moi ∼ 13600 virus particles per cell). After removal of unbound virus, incubation was continued at 37°C for additional 0, 1, 4, 7 or 11 h. Fixed cells were stained with probes against E1A and E1B-55K. Mean fluorescence intensity per cell was used to quantify the abundance of E1A and E1B-55K transcripts in cells at the different time points. Horizontal bars represent median values and the number of cells analyzed for each time point is indicated. Representative images from the time points are shown in S1 Fig. E) Correlation of E1A and E1B-55K transcripts in individual infected cells at the indicated time points post infection. The dataset is the same as in (D). r_s_ denotes the Spearman’s correlation rank coefficient (approximate P values <0.000001 for all three). F) Time course analysis of protein VI transcript accumulation. The experiment was carried out as described above for E1A and E1B-55K, except that the cells were fixed at 11, 16 and 22 h after removal of unbound virus. Cells were stained with probes against E1A and VI. The nuclear area was excluded when quantifying the abundance of transcripts, because during the late time points the probes not only marked the individual viral transcripts, but also stained the viral replication sites in the nucleus (highlighted by arrows). All images shown are maximum projections of confocal stacks. Nuclei (DAPI stain) are blue. Scale bars = 10 µm.

Under these standardized conditions, infected cells displayed high numbers of E1A transcripts at 5 h pi (Figure 1B). E1A is the first mRNA transcribed from the nuclear viral genomes. Upon early gene expression and vDNA replication, the late mRNAs for virus structural proteins, like protein VI mRNA, start to accumulate, as indicated by cell population studies [52, 58]. In accordance, protein VI transcript puncta were detected at 12 h pi, notably with high cell-to-cell variability and abundant E1A transcripts (Fig. 1B). The E1A, E1B-55K and VI signals in infected cells were specific, since the probes yielded only very occasional puncta in noninfected cells. Note that individual mRNAs appeared as distinct fluorescence puncta, but the high number of viral transcripts gives rise to clusters in the maximum projection images. RNase A treatment of the samples prior to staining removed the signals, as shown for E1A. In addition, a strong accumulation of E1A transcripts was observed in HDF-TERT infected cells albeit not in all cells and with delayed kinetics, for example at 22 h pi (Fig. 1C). HDF-TERT are nontransformed human diploid fibroblasts immortalized by human telomerase expression [5, 59]. The data indicate that the high and variable abundance of E1A transcripts is not restricted to cancer cells.

The E1A probes covered the entire E1A primary transcript region and thus all E1A splice variants. The temporal control of E1A primary transcript splicing and E1A mRNA stability give rise predominantly to 13S and 12S E1A mRNAs at 5 h pi [45, 60, 61]. The 289-residue E1A protein translated from the 13S mRNA stimulates transcription from E1A and other viral promoters, and therefore other viral early promoters are activated later than that of E1A [62]. Accordingly, although the early E1B-55K transcripts were detected in cells at the 5 h time point, these transcripts were generally less abundant than those of E1A, as indicated by representative images and quantification (Fig. 1B and 1D). Time-resolved analysis of E1A and E1B-55K transcripts was carried out in AdV-C5 infected A549 cells at moi ∼13600 virus particles per cell (37°C, 60 min incubation) yielding between 5 and 127 virus particles per cell (median 26; see Fig. 1A). Since the high abundance of viral transcripts in individual cells precluded an automated segmentation of individual transcript puncta, mean fluorescence probe intensity per cell was used to estimate viral transcript abundance. No E1A or E1B-55K transcripts were detected right after the removal of unbound virus, and only occasional cells displayed low numbers of E1A transcripts at the 2 h time point (Fig. 1D). E1A transcripts began to accumulate at 5 h pi, with high cell-to-cell variability (see also S1 Fig). Although low numbers of E1B-55K transcripts could be detected already at this time point as well, the E1B-55K transcripts began to emerge in higher numbers only at about 8 h pi. After 12 h, most cells displayed both E1A and E1B-55K transcripts. Despite considerable cell-to-cell variability, we observed good correlations in E1A and E1B-55K signals per cell at 5, 8 and 12 h pi, with Spearman’s rank correlation coefficient (r_s_) of 0.63, 0.88 and 0.79, respectively (Fig. 1E). This is in agreement with the notion that the E1B promoter is regulated by E1A [63].

The late protein VI transcripts were first detected at 12 h pi in a subset of cells, with increasing numbers of cells displaying high amounts from 17 h onwards (Fig. 1F). At the 17 h time point, about half of the cells had high numbers of protein VI transcripts, and most of them very high numbers of E1A transcripts. By 23 h, the majority of cells contained abundant VI transcripts, but the cytoplasmic E1A transcripts were reduced. Whereas other time points showed relatively few E1A, E1B-55K or VI puncta over the nuclear area, clustered nuclear E1A signals were apparent at 23 h. This nuclear E1A signal is due to binding of the E1A probe to single-stranded vDNA in the replication centers (see below). Overall, the data demonstrate that viral mRNAs accumulate in high numbers in individual infected cells, and with significant cell-to-cell variability over time.

### The E1A transcript numbers early in infection correlate moderately with the number of viral genomes per cell

AdV-C5 transcripts accumulate in high numbers in infected cells at heterogeneous rates between cells. This is especially pronounced with E1A transcripts early in infection, when occasional cells showed high accumulation of E1A mRNAs (> 100 per cell), but other cells were devoid of or showed only low numbers of E1A mRNAs. To evaluate the molecular basis of this heterogeneity, we tested whether the rate of E1A transcript accumulation correlated with the number of nuclear or cell-associated vDNA. The rationale is two-fold: first, not all cells bind equal amounts of virus (from a few to more than 170 particles / cell), and, second, due to cell-to-cell differences in the overall entry efficiency, not all incoming viruses deliver their genome into the nucleus.

We used EdC-labeled AdV-C5 (AdV-C5-EdC) and a click reaction with Alexa Fluor488-conjugated azide to visualize the number of incoming vDNAs in the cell nucleus [25]. Since interpretation of the results from vDNA-E1A mRNA correlation experiments is critically dependent on quantitative detection of viral genomes, we first determined the detection efficiency of the vDNA in infected cells at different time points pi. High percentage (∼ 88%) of the incoming EdC-labeled virus particles carried a detectable vDNA signal, as determined by anti-hexon antibody 9C12 and vDNA co-staining of virions in infected HeLa-ATCC MIB1 (Mind Bomb 1) knockout cells (S2A Fig). In these cells, incoming virus particles are targeted to the nuclear pore complex, but the capsids do not uncoat because the ubiquitin ligase activity of MIB1 is needed to trigger the disassembly [28].

We next analyzed the number of vDNA molecules in A549 cells that had been incubated with EdC-labeled AdV-C5 at 37°C for 1 h (moi ∼ 23440 virus particles per cell), and fixed 2 h or 6 h after removal of unbound virus. The number of vDNAs per total cell area or per nuclear area (defined by the DAPI-mask) was determined. The number of vDNA molecules per cell or nuclear area varied between cells at both time points (Fig. 2A). Median values for total cell-associated vDNA molecules (18.5 and 12 for 3 and 7 h samples, respectively) and nuclear vDNA (16 and 8 for 3 h and 7 h samples, respectively) indicated that there was a time-dependent reduction in the number of detected vDNA molecules. This reduction could be due to a degradation of incoming vDNAs or to a decompaction of the vDNA leading to dissipated, dim click-signals. To distinguish between these possibilities, we infected A549 cells with unlabeled AdV-C5 using similar infection conditions as with the EdC-labeled AdV-C5 and analyzed incoming vDNAs at 3 h and 7 h pi by quantitative PCR. As shown in Fig. 2B, we detected a declining trend in the vDNA numbers at 7 h pi compared to 3 h pi. This implies that the time-dependent decrease in vDNA click signals in cells infected with the EdC-labeled AdV is in part due to degradation. Thus, the clickable vDNA can be used for estimation of vDNA numbers in cells.

**Figure 2:**
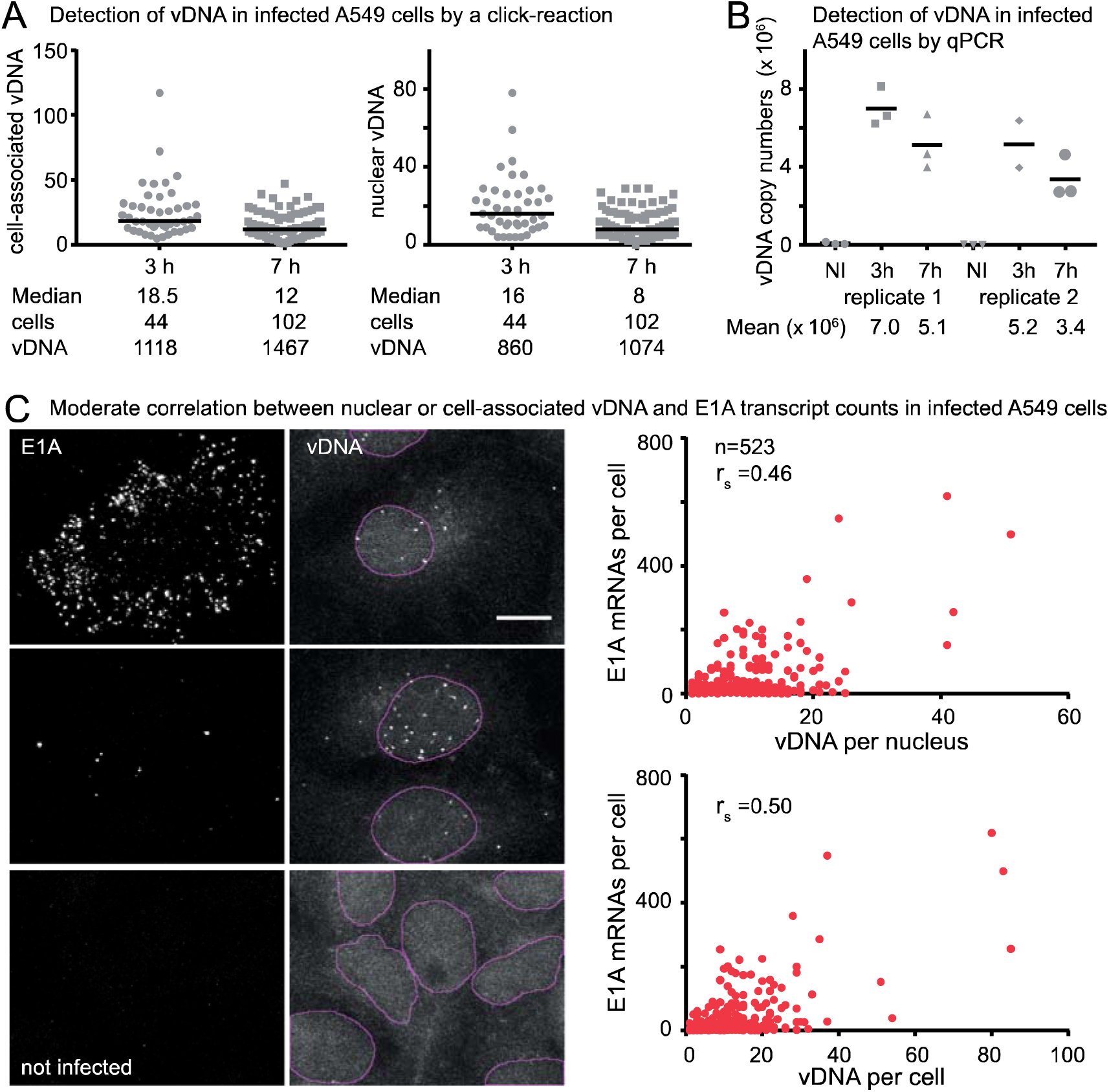
E1A mRNA abundancies at single-cell level early in infection only moderately correlate with vDNA counts in the total cell area or nucleus. A) Time-dependent decrease in the number of detected vDNAs in AdV-C5-EdC-infected cells. EdC-labeled AdV-C5 was added to A549 cells at 37°C for 60 min, and, after removal of unbound virus, incubation was continued at 37°C for additional 2 h or 6 h before fixation. The incoming viral vDNA was detected by a click-reaction using azide-Alexa Fluor488, Alexa Fluor647 NHS Ester was used for staining of cell area and nuclei were stained with DAPI. The graphs show number of vDNA molecules within the total cell area or within the nucleus area at the two time points. Horizontal lines represent median values. The number of cells and vDNAs analyzed is indicated. The differences in cell-associated or nuclear vDNA numbers between 3 h and 7 h were statistically significant (cell-associated vDNA P=0.0020 and nuclear vDNA P=0.0027, Kolmogorov-Smirnov test). B) qPCR quantification of vDNA copy numbers at 3 h and 7 h post virus addition. AdV-C5 infection conditions were similar to the experiment (A), except the virus used was not labeled with EdC. The two biological replicates and the technical replicates for each sample are shown separately. Horizontal bar represents mean. NI= noninfected control sample. C) Comparison of E1A mRNA and vDNA counts in infected cells. EdC-labeled AdV-C5 was added to A549 cells (moi ∼ 23440 virus particles per cell) at 37°C for 60 min, and after removal of unbound virus, incubation was continued at 37°C for additional 7 h. Fixed cells were stained with bDNA-FISH E1A probes and the incoming viral vDNA was detected by a click-reaction using azide-Alexa Fluor488. Alexa Fluor647 NHS Ester and DAPI stains were used for determination of cell area and nucleus area, respectively. Maximum projection images of confocal stacks were analysed using custom-programmed CellProfiler pipelines and E1A mRNA counts at single-cell level were correlated to vDNA counts per total cell area or to nuclear area. One dot represents one cell. Number of analyzed cells was 523. Since the limit for accurate segmentation of E1A puncta was about 200 per cell, counts over this number are estimates. Cells with no vDNA signal were excluded from the analysis. r_s_ denotes the Spearman’s correlation rank coefficient. Images shown are maximum projections of confocal stacks with nuclear outlines indicated. Scale bar = 10 µm.

We next addressed the question whether the number of vDNA molecules per cell or nucleus influences the rate of E1A mRNA accumulation. We incubated A549 cells with EdC-labeled AdV-C5 at 37°C for 1 h (moi ∼ 23440 virus particles per cell), removed unbound virus and incubated cells for further 7 h before fixation. Cells were stained by bDNA-FISH with E1A probes, and vDNA was visualized by a click-reaction. As shown in Fig. 2C, the E1A transcript count per cell only moderately correlated with the number of total cell-associated or nuclear vDNA puncta. A Spearman’s rank correlation test indicated r_s_ values of 0.50 and 0.46 for cell- and nucleus-associated vDNA, respectively (n=523, approximate P values <0.000001 for both). As few as five nuclear (nine cell-associated) viral genomes yielded a high number of E1A transcripts (>150 per cell), whereas several cells with 20-25 nuclear vDNA dots had only 5-39 E1A transcripts. Non-infected cells were devoid of E1A and vDNA signals. Low correlation between the number of E1A transcripts and the cytoplasmic area was observed (r_s_ value 0.23, P value <0.000001), but no significant correlation between E1A transcript numbers and nuclear area was evident (S2B Fig).

A possible population context of the cell (cell crowding, number of neighboring cells, relative position within a cell islet), which has been reported to be a dominant element in creating variable single-cell counts for cellular transcripts [64] could not be analyzed, because AdV-infected cells are less well adherent than noninfected cells and some random loss of cells was unavoidable during the RNA-FISH staining procedure.

The effect of nuclear vDNA counts on E1A transcript accumulation was analyzed in HDF-TERT cells as well. EdC-labeled AdV-C5 was incubated with the cells at 37°C for 15 h, and after removal of unbound virus, incubation was continued for additional 7 h before analysis. HDF-TERT cells are elongated cells with parts of the cytoplasmic region frequently extending over another cell. Therefore, cell segmentation had to be done manually and this explains the limited number of cells analyzed (29 cells). The result was clear though: nuclear vDNA numbers did not predict the cytoplasmic E1A mRNA counts in these cells either, for example, cells harboring ∼20 nuclear vDNAs had E1A mRNA counts ranging from 1 to 196 (S2C Fig).

### G1 cell-cycle stage promotes rapid accumulation of E1A transcripts

Results shown in Fig. 2 suggest that although the number of viral genomes per cell may contribute to the pace with which E1A transcripts accumulate early in infection, it is clear that the E1A transcript accumulation is affected by other factors as well. The cell-cycle stage has been identified as an important element of heterogeneity in cellular gene expression at single-cell level [65]. We next probed whether rapid accumulation of E1A transcripts correlated with a certain cell-cycle stage of different host cells. HeLa-FUCCI cells allow for easy classification of G1 cells (Kusabira Orange-hCdt1 expression, mKO2) and S/G2/M cells (Azami-Green-hGeminin expression, mAG) [66].

Qualitative assessment of E1A transcripts in infected HeLa-FUCCI cells at 7.5 h pi suggested that E1A transcripts accumulated more rapidly in G1 cells than in S/G2/M cells (S3A Fig). However, since the fluorescence signals of the E1A probes and mKO2 spectrally overlap, E1A transcripts over the nuclear area interfered with automated G1 vs S/G2/M classification of infected cells, and no quantitative data could be obtained from these cells. We therefore used the total intensity of the nuclear DAPI signal (a proxy for DNA content) measured by fluorescence microscopy to classify cell-cycle stages [67]. We correlated the intensities of Cdt1 and hGeminin signals in HeLa-FUCCI cells to the DNA content distribution. As shown in Fig. 3A, low DAPI intensity marked by the G1 peak correlated with high mKO2-Cdt1 nuclear signal, and high DAPI intensity with high mAG-hGeminin signal (as shown by the G2/M peak). Thus, the total intensity of nuclear DAPI signal can be used to accurately assign G1 vs S/G2/M stage to cells.

**Figure 3:**
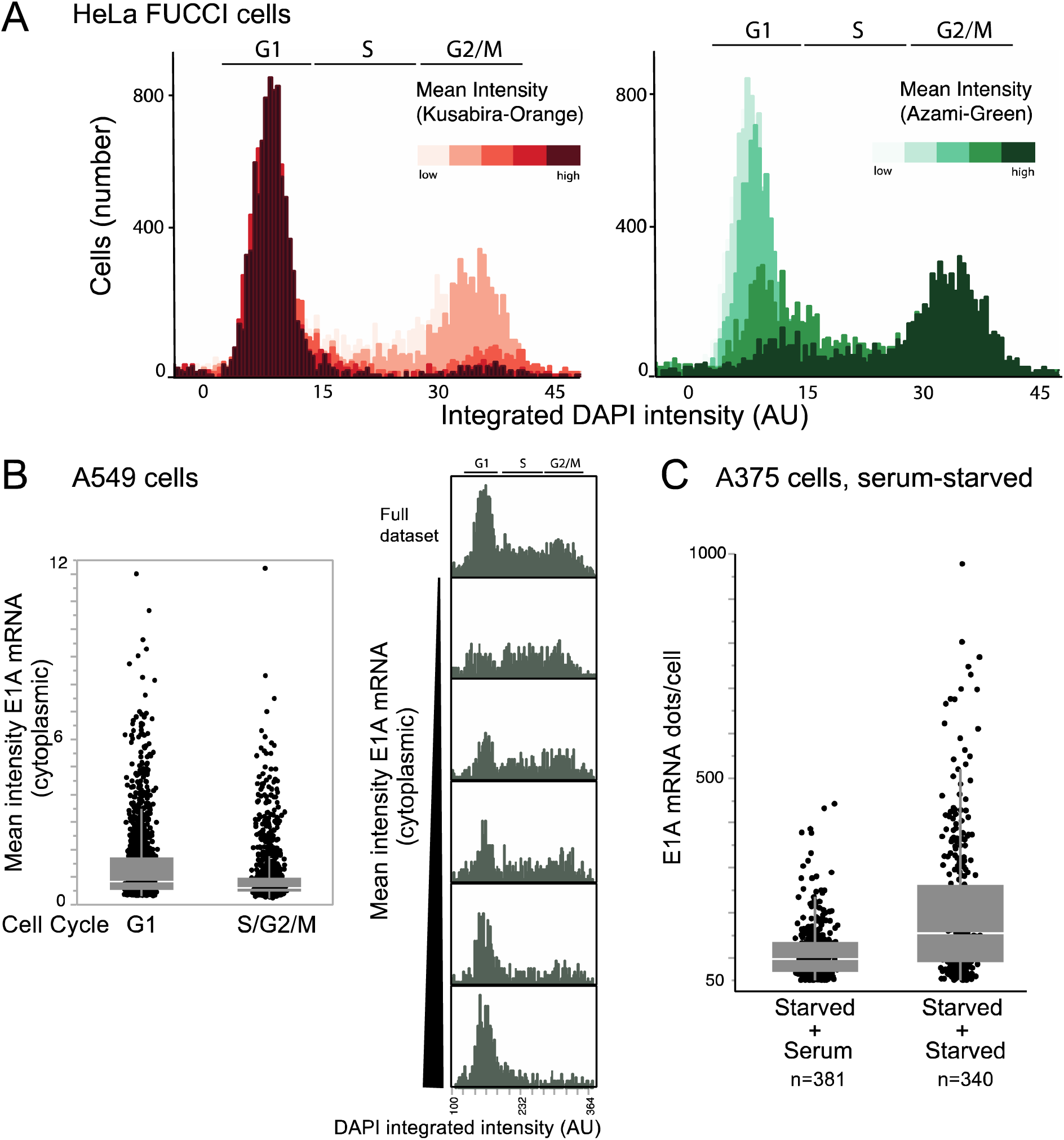
G1 cell-cycle phase favors rapid accumulation of E1A transcripts. A) Total nuclear DAPI signals can be used for accurate determination of different cell-cycle stages. Fixed noninfected HeLa-FUCCI cells were stained with DAPI and imaged by automated widefield fluorescent imaging system. The histograms show integrated nuclear DAPI intensities correlated at single-cell level to bins of increasing intensities of the G1 marker Cdt1 (Kusabira Orange) and the S/G2/M marker Geminin (Azami-Green). The increasing intensity bins of Cdt1 and Geminin are equal frequency plots in which each bin has equal number of cells. The integrated nuclear DAPI intensities are normalized to values 0-45. B) E1A transcripts accumulate more rapidly in G1 cells than in S/G2/M cells. AdV-C5 was added to A549 cells at 37°C for 60 min (moi ∼ 54400 virus particles per cell). After removal of unbound virus, incubation was continued at 37°C for additional 3 h. Fixed cells were stained with bDNA-FISH E1A probes, Alexa Fluor647 NHS Ester was used for staining of cell area and DAPI for nucleus. Images were acquired by an automated widefield fluorescent imaging system. The mean cytoplasmic E1A signal intensities were used for estimation of E1A transcript abundancies per cell. Cells were classified as G1 or S/G2/M according to total nuclear DAPI signals and 2973 cells were randomly sampled from the total population. Of these sampled cells 1659 were G1 cells and 1314 S/G2/M cells. In the left-hand panel, the mean cytoplasmic E1A signal intensities per cell in the two groups are shown as boxplots. The difference between the groups is statistically significant (permutation test, p=0.0002). In the right-hand panel, histogram of integrated nuclear DAPI intensities of cells was drawn and mean cytoplasmic intensities of E1A transcripts were mapped on the histogram. The histogram was split into five bins of increasing E1A intensities to show the correlation between E1A transcript abundancies per cell and the cell-cycle phase. Each bin contains equal number of cells. C) G1-enriched infected cell population shows increased numbers of E1A transcripts per cell in comparison to a cell population with lower number of G1 cell-cycle stage cells. AdV-C5 was added to serum-starved A375 melanoma cells at 37°C for 60 min (moi ∼ 36250 virus particles per cell), and after removal of unbound virus, cells were further incubated in serum-free (“starved + starved” sample) or serum-containing medium (“starved + serum” sample) for further 9 h. Fixed cells were stained with E1A probes, with Alexa Fluor647 NHS Ester for cell area, with DAPI for nucleus and imaged by confocal microscopy. As judged from integrated nuclear DAPI intensities, 66% of cells were in G1 phase in the “starved + starved” sample, and 42% were G1 cells in the “starved + serum” sample (S3D Fig). Cells expressing more than 50 E1A transcripts per cell were included into the data shown in the boxplot and the number of cells analyzed is indicated. Since the limit for accurate segmentation of E1A puncta was about 200 per cell, counts over this number are estimates. Permutation test indicated that the difference between the two samples is statistically significant (p=0.0002).

We initially correlated E1A transcript abundance and cell-cycle stage using a large dataset obtained by automated widefield fluorescent imaging of A549 cells incubated with AdV-C5 at 37°C for 1 h (moi ∼54400 virus particles per cell) and fixed after 3 h or 6 h following removal of unbound virus. E1A proteins induce S-phase of the cell cycle and give rise to an optimal environment for viral genome replication [reviewed in 47]. We first compared the DNA contents of noninfected and the 4 h sample of infected cells to ascertain that no viral manipulation of the cell cycle had yet taken place. Since the cell-cycle profiles and the fraction of cells in different stages of cell cycle might be affected by unequal number of cells in different conditions, we randomly sampled and selected equal number of cells from noninfected and infected samples to draw the histograms of DNA content. These histograms, as well as the visually selected cutoffs for G1 vs S/G2/M cells are shown in S3B Fig. The histograms of noninfected and infected cells were similar, and no increase of S-phase cells in the infected population was evident at this time point. The mean cytoplasmic E1A probe signal intensities were used to estimate E1A transcript abundance. When the mean cytoplasmic E1A signal intensities of the cells were split into five equal frequency bins of increasing E1A intensities, both the G1 and S/G2/M cells were found in the lowest E1A bins, whereas G1 cells clearly dominated highest E1A bin (the right-hand panel in Fig. 3B). The left-hand panel in Fig. 3B shows the distribution of E1A signals in G1 vs S/G2/M cells as a boxplot. The difference between the two classes was statistically significant according to a permutation test (p=0.0002). Furthermore, when focusing on the highest E1A expressing cells, i.e. the cells with mean cytoplasmic E1A intensities larger than 1.5 × interquartile range from the 75^th^ percentile, 71.9% of these cells were found to be in the G1 phase of cell cycle, whereas only 55.8% of cells in the total sampled cell population were G1 cells. The difference between G1 and S/G2/M cells was not an artefact of the sampling time point, since also at the 7 h time point 72.6 % of the cells with highest E1A transcript numbers were G1 cells (outliers in the S3C Fig. boxplot), whereas only 57.2 % of cells in the total sampled cell population at this time point were G1 cells. Thus, G1 cell-cycle stage favors rapid early accumulation of E1A transcripts in infected cells.

To confirm this result, we also tested E1A transcripts in infected A375 melanoma cells that were either heterogeneously distributed or enriched for G1 phase cells. The cultures were first preincubated in serum-free medium for 19 h to enrich for G1 cells, AdV-C5 (moi ∼ 36250) was added to the cells for 1 h, and, after removal of unbound virus, cells were further incubated either in the serum-free medium or switched to a serum-containing medium for additional 9 h. In the serum-starved cultures, 66% of cells were in G1 phase, whereas addition of serum reduced the fraction of G1 cells to 42% as cells rapidly moved into S-phase (S3D Fig). E1A transcript amounts per cell were determined by segmentation and counting of E1A fluorescence puncta, but since accurate segmentation was limited to about 200 E1A puncta per cell, the values above this number are only estimates. Majority (∼ 96%) of cells in both cultures had no or less than 50 E1A transcripts per cell, and these cells were excluded from further analysis and we instead focused on the remaining high E1A expressing cells. As shown by the comparison in Fig. 3C, the serum-starved, G1-enriched culture accumulated higher E1A transcript counts per cell than the serum-treated culture (p=0.0002, permutation test).

We next tested whether the more rapid accumulation of E1A transcripts in G1 cells also translates to higher E1A protein levels in this cell-cycle phase. We first analyzed EGFP signals in HeLa-ATCC cells transfected with a plasmid encoding EGFP under the control of E1A promoter and enhancer or cytomegalovirus (CMV) immediate early promoter and enhancer region. When the nuclear EGFP signal intensities in the E1A promoter-driven expression were split into four equal intensity bins of increasing intensity, the ratio of G1 vs. S/G2/M cells was different in the lowest and highest bins, with the highest bin being more clearly dominated by G1 cells (the right-hand panel in Fig. 4A). In contrast, when EGFP expression was under the control of the CMV promoter, which is most active in S-phase [68], relatively more S/G2/M cells were in the highest bin class (the left-hand panel in Fig. 4A). To test E1A protein levels, we incubated HeLa-FUCCI cells with AdV-C5 (moi ∼ 11200 for 1 h at 37°C), removed unbound virus and continued the incubation for further 9.5 h before fixation and analysis. Cells were segmented with a CellProfiler script and further sorted into early G1, G1, G1/S and S/G2/M cell-cycle stages using CellProfiler Analyst Classifier. As shown in Fig. 4B, E1A expression was most efficient not in G1, but in G1/S cells, which could reflect the advancement of high E1A expressing cells into S-phase.

**Figure 4:**
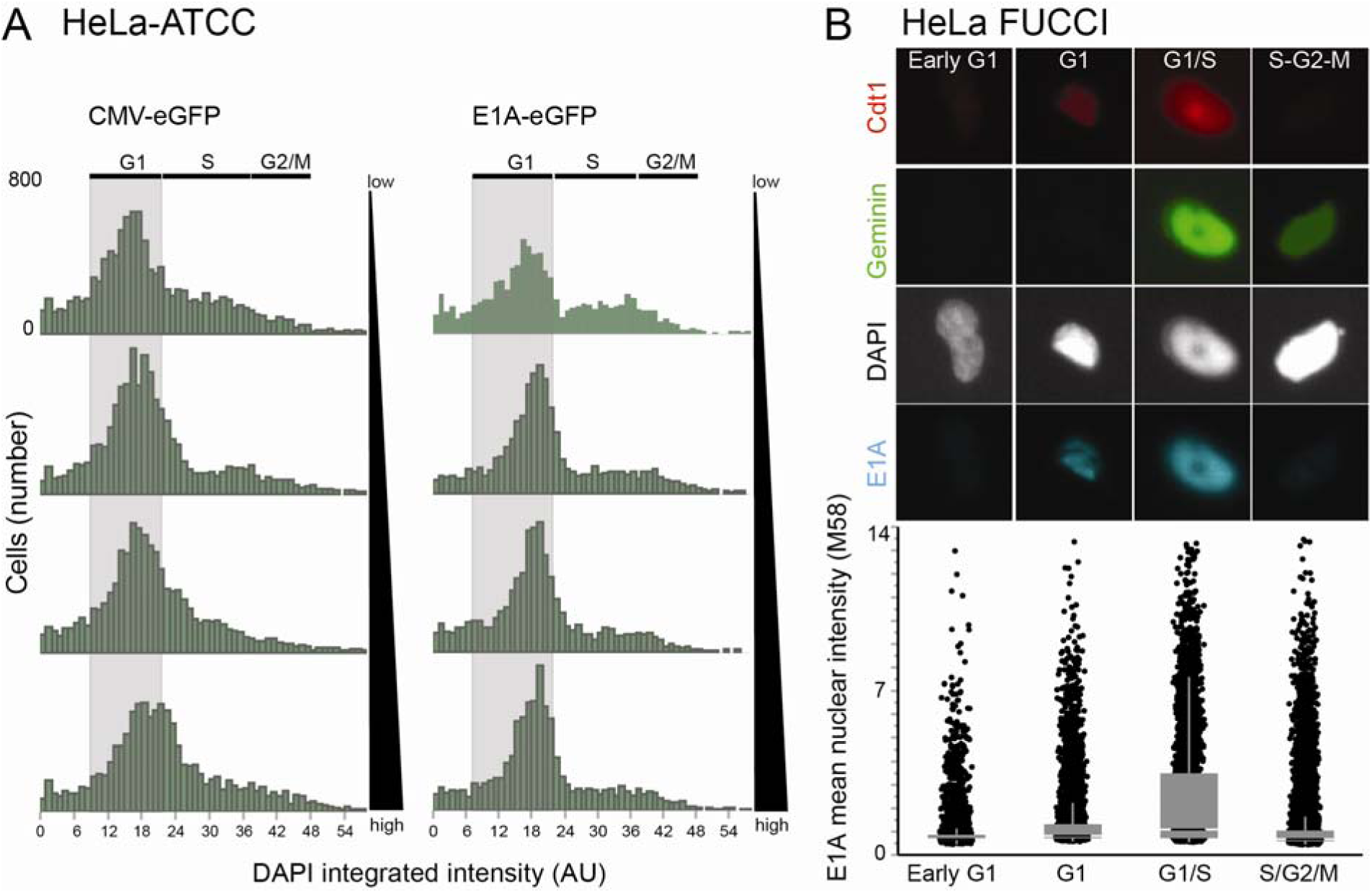
G1 and G1/S cell-cycle stages favor high E1A protein expression. A) High EGFP-expressing cells are predominantly G1 cells when the protein is expressed from E1A promoter, but in CMV promoter-driven EGFP expression S and G2/M cell-cycle phases are more favorable for high EGFP expression. The data was obtained from plasmid-transfected HeLa-ATCC cells analyzed at 48 h post transfection by automated widefield fluorescent imaging system. Histograms of integrated nuclear DAPI intensities were used to determine different cell-cycle stages and mean nuclear EGFP intensites were mapped on the histogram. The histograms represent an equal frequency plot in which mean nuclear EGFP intensities are split into four bins of increasing intensity, each bin containing equal number of cells. B) Highest E1A protein expression is seen in the G1/S cells in HeLa-FUCCI infection. HeLa-FUCCI cells were incubated with AdV-C5 (moi ∼ 11200) at 37°C for 1 h, and after removal of unbound virus cells were further incubated for 9.5 h before fixation. Cells were stained with mouse M58 anti-E1A and secondary Alexa Fluor680-conjugated anti-mouse antibodies and DAPI. Cells were imaged by automated widefield fluorescent imaging system and CellProfiler Analyst Classifier was used to assign cells into the indicated cell-cycle stages according to their Cdt1 and Geminin nuclear signals. The mean nuclear E1A intensities in the early G1, G1, G1/S and S/G2/M cells are shown as boxplots. Permutation tests with pairwise comparisons indicated that differences between G1/S and other sample populations are statistically significant (p=0.0002).

### Enhanced correlation of the E1A transcript with vDNA counts in G1 cells

Diverse conditions have been thought to contribute to variability of gene expression, including the stochastic nature of gene expression, differences in the cell-cycle state and differences in the microenvironment of cells [64, 65, 69–71]. Based on our finding that E1A mRNA abundance is cell-cycle dependent, we reanalyzed the data presented in Fig. 2 by first classifying cells into G1 vs S/G2/M (S4 Fig), and then correlating the number of cell-associated or nuclear vDNA puncta to E1A transcript numbers in the same cells. As shown in Fig. 5, the number of E1A transcripts per cell correlated better with the number of nuclear viral genomes in G1 than in S/G2/M cells. The r_s_ values were 0.52 and 0.39 for G1 and S/G2/M cells, respectively (G1 n=346 and S/G2/M n=177, approximate P values for both < 0.000001). A similar trend was observed if total cell-associated vDNA was used (r_s_ values of 0.56 and 0.41 for G1 and S/G2/M cells, respectively, P values for both < 0.000001). These results are in line with the finding that the G1 cell-cycle stage augments to the accumulation of E1A transcripts.

**Figure 5:**
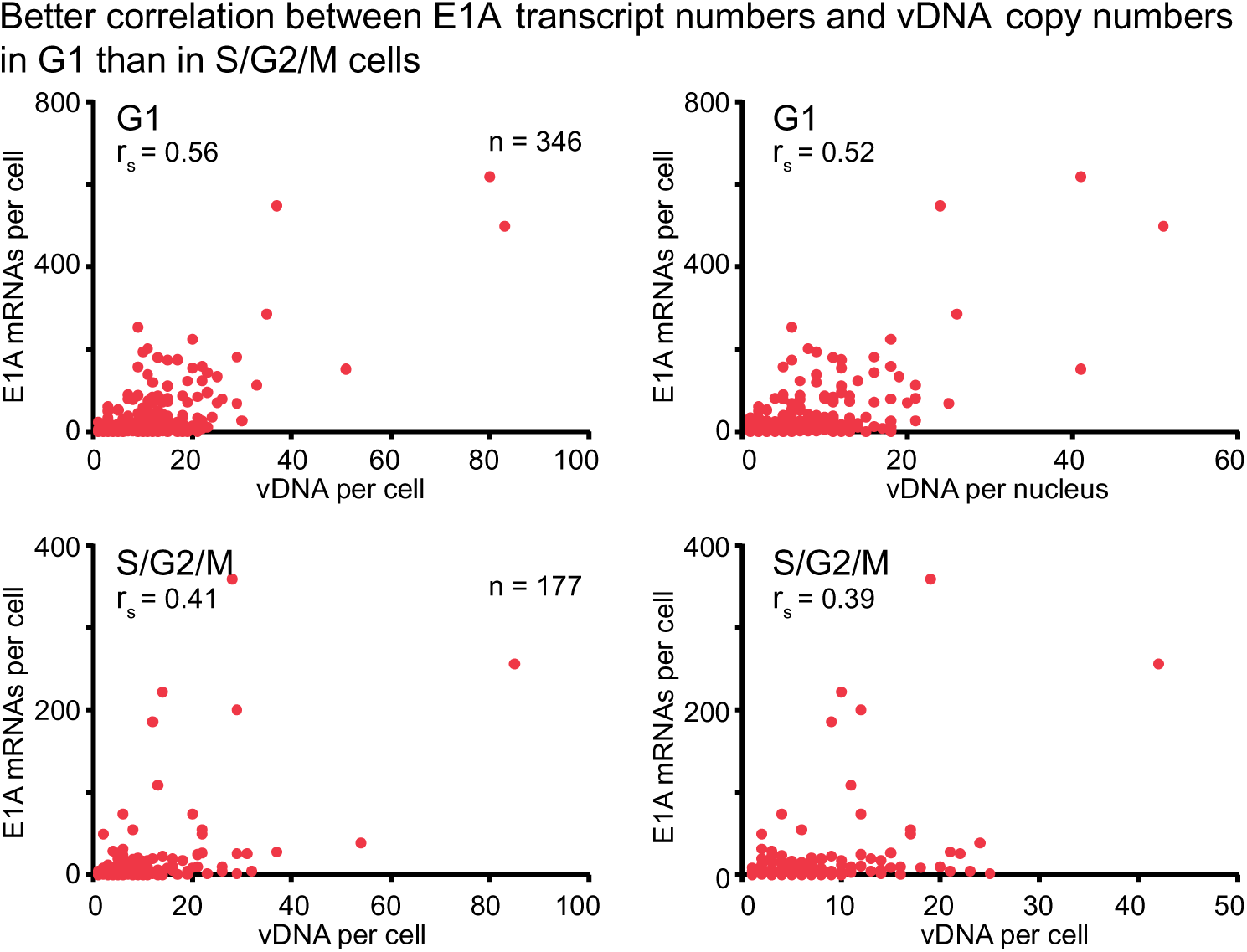
Better correlation between E1A mRNA abundancies per cell and total cell-associated or nuclear vDNA numbers in G1 cells than in S/G2/M cells. The data set is the same as in Fig. 2, but cells were first classified as G1 or S/G2/M cells according to their integrated nuclear DAPI intensities (S4 Fig) and the number of cell-associated or nuclear vDNA puncta were correlated to E1A mRNA numbers at single-cell level. One dot represents one cell. r_s_ denotes the Spearman’s correlation rank coefficient.

### High transcriptional variability of individual vDNAs in the nucleus

The enhanced correlation between the nuclear vDNA and E1A mRNA levels in the G1 cells did not explain a large fraction of cell-to-cell variability in E1A transcript amounts early in infection. We thus assessed the amount of transcriptionally active nuclear vDNA. Cellular gene loci active in transcription have been visualized by RNA FISH targeting intron sequences [72, 73], or by monitoring transcriptional bursts of nascent transcripts manifested as RNA FISH signals larger and brighter than those of individual mRNA molecules [74].

For E1A (or E1B-55K), we did not detect transcriptional bursts with bDNA-FISH probes on nuclear vDNAs, either prior to or after accumulation of viral transcripts in the cell cytoplasm. While the introns of the AdV-C5 E1A primary transcript are too short to be visualized by bDNA-FISH, the E4 primary transcript has an abundant intron of ∼ 811 bases [52]. This intron is retained in the E4orf1 and E4orf2 mRNAs. We used bDNA-FISH probes against this E4 intron and EdC-labeled incoming vDNAs to identify transcriptionally active nuclear vDNAs. A549 cells were incubated with EdC-labeled AdV-C5 at 37°C for 1 h (moi ∼ 23440 virus particles per cell) and after removal of unbound virus, incubation was continued at 37°C for additional 15 h. In contrast to E1A, E1B-55K and VI, relatively moderate numbers of E4 puncta were present in the cytoplasm, that is, a median number of 7 puncta per cell at 16 h pi, and 14 puncta at 18 h pi (S5A Fig). This is not surprising given that intronic RNAs have been estimated to be rather unstable [75].

To enhance the nuclear signals, and reduce the cytoplasmic one, we used acetic acid in the fixation buffer [56]. As shown in Fig. 6A, E4 intron probes yielded nuclear puncta of varying intensity and the brighter puncta and some of the smaller puncta colocalized with vDNA click-signals (indicated by red arrowheads). The nuclear E4 signals were mostly from RNA, and suppressed by RNase A treatment (S5B and S5C Fig). Plotting nuclear vDNA counts and the fraction of nuclear vDNAs positive for the E4 probe signal revealed cell-to-cell variability in the fraction of transcriptionally active nuclear vDNAs (Fig. 6A). In a minority of cells (9%) all nuclear vDNAs were associated with an E4 RNA and 71 % of these cells had nuclear vDNA counts ≤5 (55% of total cells analyzed had nuclear vDNA counts ≤5). Cells with nuclear vDNAs totally devoid of E4 RNA constituted 32%, and 82% of these cells had nuclear vDNA counts ≤5. In rest of the cells, a variable fraction of nuclear vDNAs contained E4 RNA and there was no obvious correlation with the number of nuclear vDNAs. Similar results were obtained at 14.5 h pi (Fig. S5D Fig).

**Figure 6:**
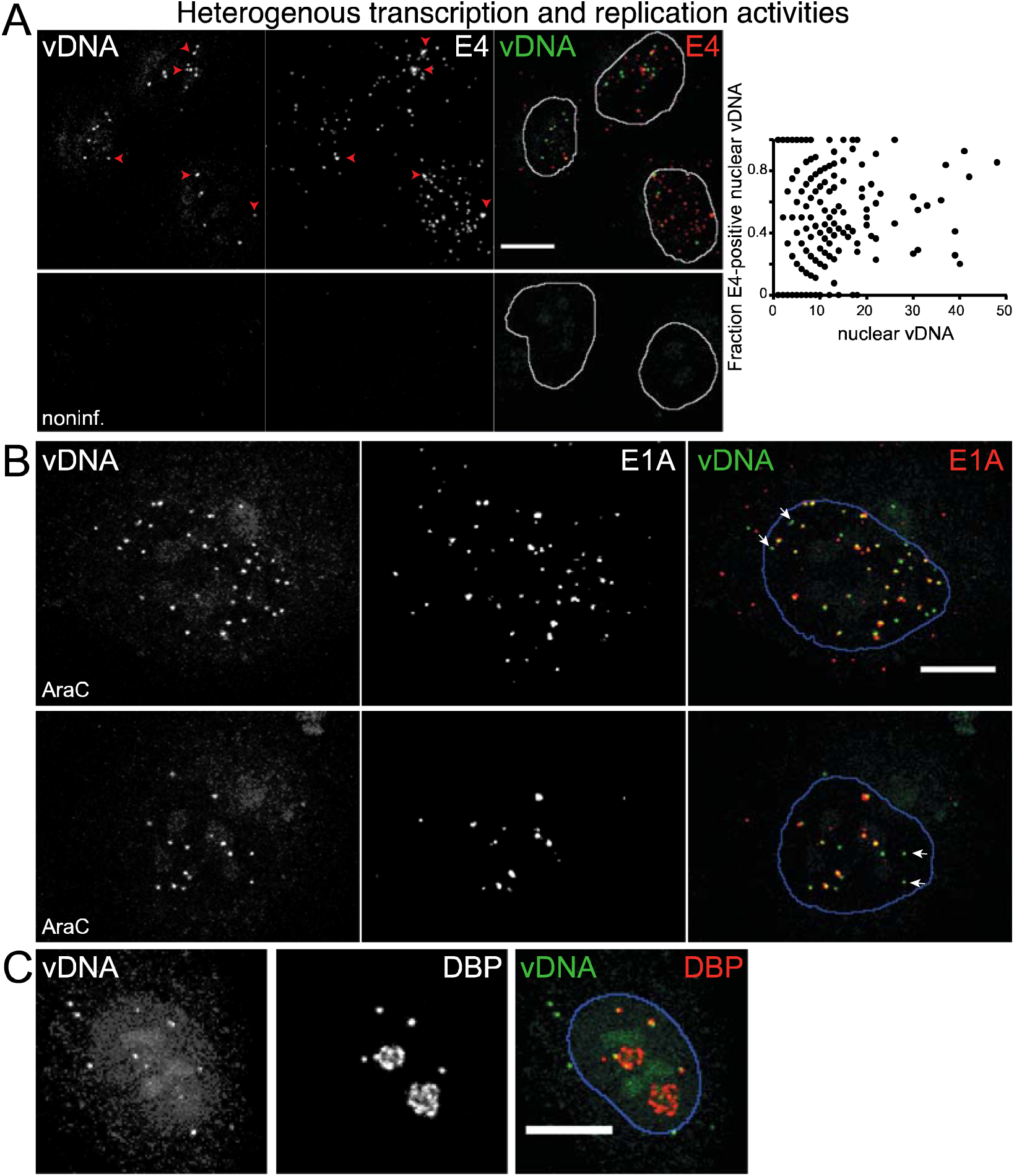
vDNAs within the same nucleus display heterogeneous transcription and replication activities. A) A549 cells were infected with EdC-labeled AdV-C5 as described in legend to Fig. 2 except that cells were fixed after 15 h post removal of unbound virus. Cells were stained with intron probes targeting the E4 transcription unit and vDNA was visualized by a click-reaction. Colocalization of nuclear vDNA with E4 probe signal indicates a transcriptionally active viral genome and the colocalization was scored from maximum projection images of confocal stacks using CellProfiler. Examples of E4 signal-positive vDNAs are indicated by red arrowheads in the representative images. Scale bar = 10 µm. In the right-hand scatterplot, nuclear vDNA numbers are correlated to the number of E4 signal-positive vDNAs within the same nucleus, one dot representing one nucleus. Number of cells analyzed was 442. Note that acetic acid was included into the fixation buffer when analyzing colocalization of E4 signals and vDNA in the nucleus, in order to improve accessibility of the E4 intron probes to nuclear targets [114]. B) Binding of E1A bDNA-FISH probes to single-stranded vDNA identifies nuclear vDNAs that have progressed to the replication stage. A549 cells were infected with EdC-labeled AdV-C5 as described in legend to Fig. 2 except that cells were fixed after 27 h post removal of unbound virus and AraC was added to the culture medium for the last 20 h. Cells were stained with E1A bDNA-FISH probes, which mark the single-stranded vDNA byproducts of viral genome replication (S6 Fig.), and the infecting vDNAs were visualized by a click-reaction. Two representative cells are shown. The majority of nuclear vDNAs at this time point post infection had progressed to the replication stage as indicated by colocalization of vDNA dots with E1A probe signals, but vDNAs devoid of E1A signal were commonly detected as well (white arrows in the overlay image). C) vDNAs within the same nucleus are associated with different stage replication centers. A549 cells were infected with EdC-labeled AdV-C5 in the absence of AraC and viral replication centers were stained with mouse anti-DBP and secondary Alexa Fluor594-conjugated anti-mouse antibodies, and incoming vDNAs were detected by a click-reaction using azide-Alexa Fluor488. Small DBP-positive puncta indicate early-phase replication centers, and both these early-phase replication centers, as well as the later-phase globular or ring-like DBP-positive structures were associated with incoming vDNAs, thus indicating that vDNAs within the same nucleus start replication asynchronously. All images shown are maximum projections of confocal stacks. Nuclei outlines were drawn from DAPI-stained nuclei. Scale bars = 10 µm.

vDNAs within the same nucleus display heterogeneous transcriptional activity at a given time point (Fig. 6A). To further analyze the varied activity status of vDNAs within a common nucleus, we analyzed the progression of incoming vDNAs to the replication phase. As shown in Fig. 1F and S6A Fig, nuclear signals with E1A probes emerged only rather late in infection and then these signals were detected within the viral replication centers. If vDNA replication was suppressed by cytarabine [AraC: S6A Fig. and 76], which creates stalled replication foci [77], a punctate nuclear pattern was observed with E1A probes. The nuclear E1A signals in AraC-treated cells were resistant to RNase A, but they were dampened by treatment with S1 nuclease (S6B Fig). Thus, these puncta, or the puncta and ring-like structures seen in cells with active vDNA replication, originate from binding of the E1A probes to single-stranded vDNA, which are a byproduct of AdV DNA replication [78].

We used this binding of E1A probes to single-stranded vDNA to monitor progression of incoming vDNAs to the replication phase. A549 cells were incubated with EdC-labeled AdV-C5 for 1 h at 37°C (moi ∼ 23400 virus particles per cell), unbound virus was removed and incubation continued for 27 h, with AraC added for the last 20 h. Representative images for colocalization of vDNA with E1A nuclear puncta are shown in Fig. 6B. The majority of vDNA in the nuclear area had an associated E1A signal, but as indicated by the white arrows in the overlay images, vDNA lacking an E1A signal could be detected as well. The vDNA puncta without E1A were present in confocal slices that contained also vDNA puncta positive for E1A signal and thus they are unlikely to originate from cytoplasmic vDNA that appeared nuclear due to a maximum projection artefact. From 53 cells analyzed, 43 (81%) had at least one nuclear vDNA dot without an associated E1A signal. Thus, individual vDNAs within the same nucleus progress to the replication phase asynchronously. This is was also seen in AdV-C5-EdC-infected cells not treated with AraC (Fig. 6C). Early stage AdV replication centers appeared as small round puncta when visualized by an antibody directed against the vDNA binding protein (DBP), and these puncta later progressed into larger globular or ring-like structures [79, 80]. The EdC-labeled viral genomes within the same nucleus could be associated with either early stage or more advanced replication centers (Fig. 6C).

## Discussion

### Transcript detection in single cells

Classical infection assays report average viral gene expression levels of a cell population. Advanced single cell technology, including RNA-FISH and single-cell RNA-Seq (scRNA-Seq) have empowered the possibility to address the viral transcription heterogeneity at the single cell level, and started to reveal significant cell-to-cell variabilty. scRNA-Seq permits simultaneous multiplexed virus and host transcriptome analysis. It has demonstrated large cell-to-cell heterogeneity in viral gene expression in influenza, Dengue and Zika flavivirus infections, and latent HIV-1 infected human primary CD4+ T cells [11, 16, 81, 82]. scRNA-Seq, however, has been technically challenging, requiring single cell isolation, RNA amplification and complex data analysis. It does not provide spatially resolved information in the infected cells which limits the interpretation of the data. For example, the finding that latent human cytomegalovirus infection differs from lytic infection only by quantitative, but not qualitative changes in the viral transcriptome has remained unexplained [83]. In addition, single transcripts containing reporter sequences, such as a hairpin structure binding to the coat protein of bacteriophage MS2, can be tracked in live cells (reviewed in [84]). Such assays are, however, invasive, and require the addition of large amounts of tandem-arranged hairpins (up to 2 kb) into the viral genome, which may interfere with the replicative ability of the virus.

To bypass all these shortcomings, we developed a novel combination of image-based single molecule assays for virus infection. We exploit the subcellular information of chemically tagged fully replicative single viral genomes, bDNA-FISH assays reporting viral transcripts and single virion fluorescence assessing the particle location. In the past, these technologies had been used individually to visualize, for example, the incoming vDNA genome, or reverse transcribed retroviral DNA [25, 85], and the bDNA-FISH technique tracked early events in HIV-1 infection [86]. Unlike scRNA-Seq, bDNA-FISH is more readily approachable, and provides subcellular information. For example, multiple single labeled RNA-FISH probes monitored lymphocytic choriomeningitis mammarenavirus RNA species during acute and persistant infections [12], or probed the genome segment ratios and the packaging mechanism of the tri-segmented bunyavirus Rift Valley fever virus genome in infected cells [87].

Single-molecule RNA-FISH assays are convenient to use, since assay systems are commercially available (ViewRNA, RNAscope, Stellaris), RNA molecules are detected without isolation or enzymatic amplification procedures with single-molecule and subcellular resolution, and data analysis is straightforward with open-source tools, such as CellProfiler and Knime. However, the assays also have drawbacks, for example, a relatively long target sequence is required for a bright signal, which makes mRNA splice variant detection challenging. Another drawback for RNA-FISH technologies in general is the limited number of targets that can be simultaneously uncovered, although different schemes for sequential rounds of hybridization have been developed to multiplex single-cell RNA-FISH [88, 89]. Yet, such procedures are not necessarily applicable for virus infections if cells do not remain firmly attached to imaging plates, as in the case of AdV-C5.

The bDNA technology and click chemistry developed in this study allows for quantitative direct detection of mRNAs at single-molecule level and enables the monitoring of virus transcript accumulation from entry to replication and egress. Using this approach, we relate the rate of accumulation of the first viral mRNA, the E1A mRNA, to the number of viral genomes. The data show that in some cell nuclei but not others, the vDNAs have non-uniform transcriptional activity and progress asynchronously into the replication phase. Specifically, our results indicate that the E1A, E1B-55K and VI viral transcripts accumulate in high numbers in infected cancer cells and non-transformed cells, such as HDF-TERT cells, typically reaching levels >200 transcripts per cell, in agreement with population-based assays [42, 58, 90]. In comparison, only about 5% of HeLa cell mRNAs have mean transcript numbers larger than 200 per cell [56]. Of note, however, the overall infection kinetics of HDF-TERT cells were slower than in the A549 cancer cell, but again, no clear correlation between E1A transcript and nuclear vDNA numbers was observed across the population.

### Low correlation of viral genome and transcript abundance

Our studies underscore that viral genomes within a single nucleus can give rise to variable amounts of viral mRNA transcripts. For example, similar high amounts of E1A mRNAs (>100) were observed in cells with less than ten or more than 40 nuclear vDNAs. Conversely, examples of cells with E1A transcripts less than 20 were easily found in cells with nuclear vDNA counts of ∼20, although in general, cells with high nuclear vDNA counts (>40) contained high E1A transcript numbers (> 100) at 8 h pi. At present, the molecular basis of the variability between viral genomes and E1A expression early in infection is unknown. Transcription from the E1A promoter is subject to both positive and negative regulation by cellular factors, and, in addition, E1A proteins, primarily the long 289-residue isoform, increase E1A transcription through a positive feedback loop [5, 47, 63, 91]. Abundance and intracellular localization of the cellular regulators of E1A transcription could influence the rate, at which individual cells accumulate E1A transcripts. This suggests that effective local concentrations of transcription regulators within the immediate vicinity of the vDNA rather than the overall cellular or nuclear concentrations will determine the transcriptional output from the viral promoters. Indeed, several reports have provided evidence that transient, dynamic spatio-temporal clustering of transcription regulatory factors and RNA polymerase II play an important role in transcription output from cellular promoters [92–98].

### Enhanced E1A transcript abundance in G1 cells

What has also become clear from our study is that the cycle phase accounts for some of the cell-to-cell variability in AdV transcription. Manipulation of the cell cycle is common in infection [99]. For example, the viral E1A proteins together with key cellular interaction partners are well known to push the host cell into S-phase for optimal support of viral genome replication [47, 100]. Here we show that G1 cells promoted the rapid accumulation of E1A transcripts. The underlying reasons are currently unknown, but could involve enhanced viral transcription or decreased E1A mRNA decay, the latter reported to be cell type dependent [101]. Enhanced transcription could be mediated by positive transcription elongation factor beta (P-TEFb), which is known to enhance E1A expression, most prominent in G1, and requires BRD4, a reader of acetylated histones [102, 103]. Another mechanism could involve putative SP1 transcription factor binding sites in the E1A enhancer/promoter, as the SP1 transcription factor is expressed more in G1 phase [104].

### Variable transcription and replication activities of viral genomes

Individual vDNA molecules within the same nucleus did not show homogenous transcriptional activity, when transcriptional activity was measured by colocalization of EdC-labeled vDNA click-signal with a probe against an intron from the viral E4 transcription unit, the only early viral intron that is long enough and abundant enough to be detected by the RNA FISH procedure. This gives a snapshot view of the E4 promoter activity, and it does not reveal whether the inactive vDNAs exhibit transient or prolonged transcriptional inactivity, or whether they would transcribe other viral genes. Live assays with uninfected cells have shown that transcription of cellular genes is a discontinuous process, with periods of transcription bursts followed by inactive periods of variable length [69, 71]. This is in agreement with local transient clustering of transcription regulatory factors and RNA polymerase II. If AdV transcription followed a similar pattern and if the transcription silent periods were short, then our snapshot assay might overestimate the intranuclear heterogeneity of vDNA transcription activities.

However, the viral genomes in the nucleus not only showed transcriptional but also high replicative variability, as indicated by the progression of individual EdC-labeled incoming vDNAs into replicative centers containing ssDNA, a byproduct of viral genome replication. This emphasizes that vDNAs within the same nucleus are subjected to a differential regulation, in agreement with single cell analyses of vDNA and viral transcripts by a padlock probe (PLP)-based rolling circle amplification (RCA) used in conjunction with fluorescence microscopy, showing considerable heterogeneity in vDNA and viral mRNA contents between cells upon replication [105]. Unfortunately, the PLP-based RCA has limited sensitivity and cannot assess individual mRNAs and vDNAs.

Overall, it is likely that the three-dimensional host genome architecture around the vDNA affects the transcriptional output from the viral genome. This notion is in line with a recent Hi-C and vDNA capture analysis of AdV-C5-infected human hepatocytes where the viral genome preferentially interacted with transcription start sites and enhancers of active cellular genes, including genes that are upregulated during infection [106]. Since the Hi-C measurements are population-averaged snapshots and do not give information on the dynamics of the contacts, it is unknown whether AdV usurps enhancers of cellular genes to promote its transcription in addition to its own enhancers, or whether the viral enhancers create transcriptional hubs that benefit nearby cellular genes. Such questions can now be addressed with image-based single cell, single molecule assays described here.

In summary, our study demonstrates that the progression of AdV infection is variable at the single-cell and single genome levels, an observation that may apply to many other infections [10–16]. The strategy introduced in this study is highly versatile, and can be easily adapted to other viruses, non-viral transcripts or transgenes from viral vectors at the single-cell level in different cell types. We envision further technical developments, such as assays for live measurements of incoming viral genomes, transcriptional output and transcription regulatory factor dynamics at single viral genomes in the nucleus [107]. Improvements in single molecule detection methods [97] might make these assays feasible within near future.

## Supporting information

Key resources

## Acknowledgments

We thank Patrick Hearing, Kathleen Rundell, Laurence Fayadat, Wiebe Olijve, Cornel Fraefel and Alex Hajnal for cell lines, Nancy Reich for antibodies, Nicole Meili and Melanie Grove for tissue culture and the ZMB microscopy and image analysis core facility at the University of Zurich for advice with confocal microscopy. This work was supported by grants from the Swiss National Science Foundation (31003A_179256 / 1, 316030_170799 / 1, and CRSII5_170929 / 1) and the Kanton Zurich.

## Authors contributions

Conceptualisation (MS, VP, UFG); coordination (UFG), formal analyses and validations, data curation, software, design and conduction of cell-based experiments (MS, VP); data analyses and interpretation (MS, VP, AK, UFG); first draft of manuscript and figures (MS) with input from VP; final draft of manuscript and figures (MS, UFG) with input from VP and AK, funding acquisition (UFG).

## Materials and methods

### Cells

Two different clones of human lung epithelial carcinoma A549 cells were used in the study: our laboratory’s old A549 clone (experiments shown in Fig. 1, Fig. 3B and S1 Fig., S3B and S3C Fig., S6A and S6B Fig., RNase A treatment) and A549 from American Type Culture Collection (ATCC, experiments shown in Fig. 2 and Fig. 5, Fig. 6, S2B Fig., S4 Fig., S5 Fig., and S6B Fig. S1 nuclease-treatment). Highly-polymorphic short tandem repeat loci profiling indicated ∼ 95.1% similarity for these two A549 clones. A549 cells were maintained in DMEM (Sigma-Aldrich, D6429) supplemented with 7.5% fetal calf serum (FCS; Gibco/ThermoFisher Scientific, 10270106) and 1% nonessential amino acids (Sigma-Aldrich, M7145). HeLa cells were from ATCC (clone CCL-2) and HeLa-FUCCI cells, kindly provided by Cornel Fraefel, were originally obtained from Atsushi Miyawaki [66]. HeLa-ATCC-MIB1 knockout cells have been described before [28]. Immortalized human diploid fibroblast HDF-TERT cells expressing the catalytic subunit of telomerase were kindly provided by Patrick Hearing and Kathleen Rundell [59]. Melanoma A375 cells, kindly provided by Alex Hajnal, were originally from ATCC (CRL-1619). HeLa and HeLa-FUCCI cells, as well as HDF-TERT and A375 cells were maintained in DMEM supplemented with 10% FCS and 1% nonessential amino acids.

### Viruses

AdV-C5 viruses were grown in A549 cells and purified on CsCl gradients as previously described [108]. EdC-labeled AdV-C5 was produced in A549 in presence of 2.5 µM EdC added at 14 h post infection [25]. EdC was from Sigma-Aldrich (T511307). Absorbance measurements at 260 nm were used for determination of number of virus particles per ml [109].

### Virus binding to cells

A549 cells grown on 96-well imaging plate (Greiner Bio-One, 655090) were incubated with AdV-C5 (moi ∼ 54400 or 13600 virus particles per cell) at 37°C for 60 min in DMEM supplemented with 0.2% bovine serum albumin (BSA, Sigma-Aldrich, A9418) and penicillin-streptomycin (Sigma-Aldrich, P0781, final concentration penicillin 100 units per ml and streptomycin 0.1 mg per ml). After removal of unbound virus, incubation was continued in DMEM supplemented with 7.5% FCS, 1% nonessential amino acids and penicillin-streptomycin for 5 min at 37°C. Cells were placed on ice and stained with Alexa Fluor647-conjugated wheat germ agglutinin (ThermoFisher Scientific, W32466, 4 µg/ml in cold RPMI-1640 medium, Sigma R7388) for 1 h in dark for cell outlines. Cells were subsequently fixed with 3% paraformaldehyde in phosphate-buffered saline (PFA-PBS) for 20 min at room temperature (RT) and stained with mouse 9C12 anti-hexon antibody [110]; 9C12 antibody, developed by Laurence Fayadat and Wiebe Olijve, was obtained from Developmental Studies Hybridoma Bank developed under the auspices of the National Institute of Child Health and Human Development), secondary Alexa Fluor488-conjugated goat anti-mouse (ThermoFisher Scientific, A11029, 4 µg/ml) antibodies and 1 µg/ml DAPI (4’,6-diamidino-2-phenylindole) as described in [21]. Cells were imaged in PBS with a Leica SP5 confocal laser scanning microscope using 63× magnification oil objective (numerical aperture 1.4) and zoom factor 2. Single slice through the middle of the cell was recorded for DAPI-channel and stacks were recorded at 0.5 µm intervals for the 9C12 signal with 4× frame averaging. Number of cell-associated virus particles were determined from maximum projections of confocal stacks using a custom-programmed MatLab (The Mathworks) routine. Scatterplots were made with GraphPad Prism (GraphPad Software, La Jolla, CA, USA).

### RNA FISH with bDNA signal amplification

A549 cells were seeded at a density of 6000 per well in a 96-well imaging plate (Greiner Bio-One, 655090) and grown for two days. AdV-C5 was incubated with cells at the indicated multiplicities of infection (moi; two cell doubling times were included into the moi calculations) at 37°C for 60 min in DMEM supplemented with 0.2% BSA and penicillin-streptomycin. After removal of unbound virus, incubation was continued in DMEM supplemented with 7.5% FCS, 1% nonessential amino acids and penicillin-streptomycin for the indicated times. Cells were fixed with 3% PFA-PBS for 30 min at room temperature (RT), washed twice with PBS, dehydrated by sequential incubations in 50% ethanol for 2 min, 70% ethanol for 2 min, 100% ethanol for 2 min and plates were stored in 100% ethanol in −20°C until proceeding with RNA FISH staining. Affymetrix QuantiGene ViewRNA HC screening assay system (available from ThermoFisher Scientific) was used for single molecule RNA FISH with bDNA signal amplification [56, 57]. Cells were rehydrated by sequential incubations in 70% ethanol for 2 min, 50% ethanol for 2 min and PBS for 10 min, permeabilized by a 5-min incubation in 0.2% Triton X-100 in PBS and washed twice in PBS. RNA FISH with bDNA signal amplification was performed according to a protocol recommended by the manufacturer using custom-made probes against AdV-C5 E1A mRNAs (type 1 probes, Alexa Fluor546, probes were made against the sequence between the AdV-C5 genome map positions 551-1630), E1B-55K mRNA (type 4 probes, label Alexa Fluor488, probes were made against the sequence spanning the AdV-C5 map positions 2421-3495), VI mRNA (type 4 probes, label Alexa Fluor488, probes were made against the sequence spanning the AdV-C5 map positions 18040-18700) and E4orf1/orf2 mRNAs/introns (type 1 probes, label Alexa Fluor546, probes were made against the E4 transcript region spanning the AdV-C5 map positions 35547-34735). Nuclei were stained with DAPI (1 µg/ml in PBS) for 20 min at RT and Alexa Fluor647 NHS ester (A20006, ThermoFisher Scientific, 0.5 µg/ml in PBS for 10 min) was used for staining of the cell area. To confirm that the cytoplasmic probe signals represent viral transcripts, samples were treated or not with 65 µl of 50 µg/ml RNase A in PBS per well for 30 min at RT after permeabilization with Triton X-100, washed three times with PBS and, prior to proceeding to the RNA FISH staining, cells were again fixed with 3% paraformaldehyde in PBS for 15 min at RT. Imaging was carried out with a Leica SP5 confocal laser scanning microscope (Fig. 1 experiments) using 63× magnification oil objective (numerical aperture 1.4) and zoom factor 2. Stacks were recorded at 0.5 µm intervals with sequential acquisition using between frames switching mode and typically 4× frame averaging for the RNA FISH signals. Representative images shown are maximum projections of confocal stacks and images were processed with Fiji [111], applying the same changes in brightness and contrast to all image groups in the series. Custom-programmed CellProfiler (http://cellprofiler.org, [112]) pipelines were used to score mean cell-associated transcript intensities at single-cell level from maximum projections of image stacks. The resulting data were sorted using KNIME Analytics Platform (https://www.knime.com/knime-software/knime-analytics-platform). GraphPad Prism was used for creating the scatterplots.

For analyzing E1A transcripts in infected HDF-TERT, cells were seeded at a density of 8000 per well in a 96-well imaging plate and grown for two days. AdV-C5 was incubated with the cells at 37°C for 12 h (moi ∼ 37500 virus particles per cell) in DMEM supplemented with 10% FCS, 1% nonessential amino acids and penicillin-streptomycin. After removal of the inoculum medium, incubation was continued at 37°C for additional 10 h before cells were fixed and stained for E1A mRNAs and cell area as described above. Imaging was carried out with a Leica SP5 confocal laser scanning microscope as described above.

### Detection efficiency of incoming EdC-labeled vDNA

To determine the number of EdC-labeled virus particles carrying a detectable vDNA signal, HeLa-ATCC-MIB1 knockout cells were incubated with EdC-labeled AdV-C5 (moi ∼ 23440 virus particles per cell) at 37°C for 60 min in DMEM supplemented with 0.2% BSA and penicillin-streptomycin, and, after removal of unbound virus, incubation was continued at 37°C for 60 min in DMEM supplemented with 7.5% FCS, 1% nonessential amino acids and penicillin-treptomycin before fixation. To tag virus particles, fixed cells were stained with 9C12 anti-hexon and secondary Alexa Fluor594-conjugated anti-mouse (ThermoFisher Scientific, A21203) antibodies as described above and in [21]. The viral vDNA was detected by a click-reaction using Alexa Fluor488-conjugated azide (ThermoFisher Scientific, A10266) as previously described [25], and nuclei were stained with DAPI. The samples were imaged with a Leica SP5 confocal laser scanning microscope using 63× magnification oil objective (numerical aperture 1.4) and zoom factor 2. Stacks were recorded for all channels at 0.5 µm intervals with sequential acquisition using between frames switching mode and 3× frame averaging for the 9C12 signals and 4× frame averaging for vDNA signals. The sensitive Leica HyD hybrid detector was required for proper detection of the vDNA signals. A custom-programmed MatLab (The Mathworks) routine was used to determine the number of cell-associated virus particles from maximum projections of confocal stacks and to score vDNA signal on the virus particles. The threshold value for a positive vDNA signal was determined by placing a virus image on a click-reaction image obtained from cells infected with not EdC-labeled virus and taking the highest virus-associated signal as a cutoff value.

To determine the efficiency of incoming vDNA detection in relation to increasing infection time, A549 cells were incubated with EdC-labeled AdV-C5 (moi ∼ 23440) at 37°C for 60 min as described above, and, after removal of unbound virus, incubation was continued at 37°C for additional 2 or 6 h before fixation. The incoming viral vDNA was detected by a click-reaction using azide-Alexa Fluor488, Alexa Fluor647 NHS Ester was used for staining of cell area and nuclei were stained with DAPI. The samples were imaged with a Leica SP5 confocal laser scanning microscope using 63× magnification oil objective (numerical aperture 1.4) and zoom factor 2. Stacks were recorded at 1 µm intervals with sequential acquisition using “between stacks” switching mode. Signals for vDNA were recorded with 6 times frame averaging and with a Leica HyD hybrid detector in the normal mode. A custom-programmed CellProfiler pipeline was used to score vDNA puncta within the total cell area (Alexa Fluor647 mask) and within the nuclear area (DAPI mask). To improve vDNA puncta segmentation, vDNA images were processed with the Fiji built-in plugin Rolling Ball Background Subtraction with rolling ball radius set to 20 pixels prior to running the CellProfiler pipeline. The resulting data were sorted using KNIME Analytics Platform and GraphPad Prism was used for creating the scatter plots.

### qPCR quantification of vDNA in infected cells

Confluent A549 cell cultures on 6-well dish were incubated with AdV-C5 (moi ∼ 17600) at 37°C for 60 min as described above, and, after removal of unbound virus, incubation was continued at 37°C for additional 2 or 6 h. Noninfected cells were used as a control. Cells were washed twice with PBS, scraped into 200 µl PBS, and total DNA was extracted using Qiagen DNeasy Blood and Tissue kit (69506) according to a protocol recommended by the manufacturer. The DNA was eluted into 100 µl of the kit buffer AE. Quantification of the vDNA was performed using quantitative PCR ABI QuantStudio 3 Real-Time PCR system by setting up a three-step melt-curve analysis using primers E1A_forward 5’-GGTGGAGTTTGTGACGTGG-3’, E1A_reverse 5’-CGCGCGAAAATTGTCACTTC-3’ against the Ad5 E1A promoter and enhancer DNA [91]. Viral genome copy numbers were estimated from a plasmid standard curve.

### Relating E1A transcripts to vDNA at single-cell level

A549 cells were seeded at a density of 40000 on alcian blue-coated coverslips in a 24-well plate format and grown for two days. EdC-labeled AdV-C5 was incubated with cells (moi ∼ 23440 virus particles per cell) at 37°C for 60 min in DMEM supplemented with 0.2% BSA and penicillin-streptomycin. After removal of unbound virus, incubation was continued in DMEM supplemented with 7.5% FCS, 1% nonessential amino acids and penicillin-streptomycin for further 7 h before fixation. RNA FISH with E1A probes was carried out as described above. Coverslips were subsequently inverted on 30 µl droplets of ImageiT FX Signal Enhancer (ThermoFisher Scientific, I36933) and incubated at RT for 30 min. After two washes with PBS, click-reaction with Alexa Fluor488-conjugated azide was performed as described in [25]. Nuclei were stained with DAPI and cell area with Alexa Fluor647 NHS ester. Imaging was carried out with a Leica SP5 confocal laser scanning microscope using 63× magnification oil objective (numerical aperture 1.4) and zoom factor 2.5. Stacks were recorded at 1 µm intervals with sequential acquisition using between stacks switching mode and 3 or 6 times frame averaging for E1A and vDNA signals, respectively. The sensitive Leica HyD hybrid detector in the normal mode was required for proper detection of the vDNA signals. Maximum projections of confocal stacks and a custom-programmed CellProfiler pipeline were used to determine the E1A transcript numbers in the cell area (Alexa Fluor647 cell mask) and vDNA numbers in the total cell area (Alexa Fluor647 cell mask) or the nuclear area (DAPI mask). Proper cell and nucleus segmentation was controlled and adjusted manually if necessary. The resulting data were sorted using KNIME Analytics Platform and cells with no vDNA signal were excluded from the analysis. GraphPad Prism was used for producing the scatter plots and performing Spearman’s correlation tests. For determining the effect of cell-cycle stage for vDNA-E1A transcript number correlations, cells were first classified as G1 or S/G2/M cells according to their integrated nuclear DAPI intensities (see below) and the two cell populations were separately analyzed in GraphPad Prism for correlations between total cell-associated or nuclear vDNA counts and the number of E1A transcripts per cell. Representative images shown in figures are maximum projections of confocal stacks and images were processed with Fiji, applying the same changes in brightness and contrast to all image groups in the series.

For the HDF-TERT experiment, cells were incubated with EdC-labeled AdV-C5 at 37°C for 15 h in DMEM supplemented with 10% FCS, 1% nonessential amino acids and penicillin-streptomycin. After removal of the inoculum medium, incubation was continued at 37°C for additional 7 h before cells were fixed and processed as described above for the A549 infection.

### Assigning cell-cycle phase from integrated DAPI intensities

For determination of the cell-cycle stage, cells were stained with DAPI and imaging was performed with either wide-field high-throughput or confocal microscopy. The segmentation of the nucleus was done with CellProfiler pipeline and the DAPI intensities were measured over the nuclear mask. Following this, histograms of integrated DAPI intensities were plotted for infected and non-infected samples in statistical software JMP (JMP^®^, Version 13, SAS Institute Inc., Cary, NC, 1989-2007). The cells were called as G1 stage cells if their integrated DAPI intensities fell between the range determined by the visually selected cutoffs from the non-infected samples, as has been described before [67]. For example, in the S3B Fig., threshold for G1 population cells was between 132-200 AU (arbitrary units). Cells outside this range were called S/G2/M phase cells.

### Effect of cell-cycle on early accumulation of E1A transcripts

A549 cells grown on 96-well imaging plate were incubated with AdV-C5 at 37°C for 60 min (moi ∼ 54400 virus particles per cell) in DMEM supplemented with 0.2% BSA and penicillin-streptomycin. After removal of unbound virus, incubation was continued at 37°C for additional 3 or 6 h in DMEM supplemented with 7.5% FCS, 1% nonessential amino acids and penicillin-streptomycin. Fixed cells were stained with E1A probes, Alexa Fluor647 NHS Ester was used for staining of the cell area and DAPI for nucleus as described above. Images were acquired with Molecular Devices automated ImageXpress Micro XL widefield imaging system using 20× S Fluor objective (numerical aperture 0.75), stacks for RNA FISH and cell area channels, and single focal plane for DAPI. A custom-programmed CellProfiler pipeline was used for determining the nuclear DAPI intensities and the mean E1A transcript intensities at single-cell level from maximum projections of image stacks. The separation of cell-cycle stages between G1 and S/G2/M phases was done as described above, with threshold of 132-200 AU. The histogram in Fig. 3B (distribution of E1A mRNA cytoplasmic intensities in G1 vs S vs G2M) was drawn from a full dataset, whereas the Fig. 3B scatterplot was drawn from a randomly sampled populations of 1659 and 1314 G1 and S/G2/M infected cells, respectively. Both the histogram and the scatter plot were made using JMP. The outliers of the E1A expressing population were the cells having intensities more than 1.5× interquartile range from the 75^th^ percentile.

Analysis of early E1A transcript accumulation in G1-enriched cell population was performed in the melanoma A375 cells since these cells respond well to serum starvation. Cells were seeded on 96-well imaging plates at a density of 18000 and after one day incubation cells were switched to plain DMEM medium without FCS. After 19 h, AdV-C5 was added to cells at moi of ∼ 36250 virus particles per cell for 60 min at 37°C in plain DMEM supplemented with penicillin-streptomycin. After removal of unbound virus, incubation was continued at 37°C for further 9 h either in plain DMEM-penicillin-streptomycin (starved + starved sample) or in DMEM supplemented with 7.5% FCS, 1% nonessential amino acids and penicillin-streptomycin (starved + serum sample). Cells were fixed and stained with E1A transcript probes and DAPI as described above. For determination of E1A transcript numbers per cell, Molecular Devices automated ImageXpress Micro confocal imaging system and 40× Plan Apo Lambda objective (numerical aperture 0.95) was used for image acquisition with confocal stacks for E1A channel, and single focal plane for DAPI and a transmission light images. For determination of the percentage of G1 cell-cycle phase cells in the different samples, Molecular Devices automated ImageXpress Micro XL widefield imaging system and 10× Plan Fluor objective (numerical aperture 0.3) was used for recording the images in DAPI channel. A custom-programmed CellProfiler pipeline was used for determining the nuclear DAPI intensities and the number of E1A transcripts per cell from maximum projection image stacks (cell boundaries segmented from the transmission light images). The saved E1A and cell outline segmentation images from the CellProfiler pipeline were used to control proper segmentation and cells erroneously having numerous mRNAs as a result of a spillover from a neighboring high expressing cell were manually removed from the dataset. Only cells with more than 50 E1A transcript dots were included into the data analysis and boxplots from these cells were drawn using JMP. To check the efficiency of G1 accumulation by serum-starvation, histograms of equal numbers of randomly sampled cells of indicated samples were plotted using their integrated DAPI intensity. Cells were scored as G1-phase cells if their integrated DNA intensity fell between the visually selected cutoff of 2-15 AU and subsequently percentage of G1 phase cells was calculated.

### Comparison of effect of cell-cycle on CMV immediate early and E1A promoter activities

HeLa-ATCC cells were seeded on 96-well imaging plate at a density of 10000 cells/well and grown for one day. Plasmids containing EGFP expression cassette under the control of AdV-C5 E1A or cytomegalovirus (CMV) promoter and enhancer regions were transfected at an amount of 100 ng / well using Lipofectamine 2000 transfection method (ThermoFisher Scientific, 11668019). Cells were fixed 48 h post transfection, stained with DAPI and imaged with Molecular Devices automated ImageXpress Micro XL widefield imaging system using 20× S Fluor objective (numerical aperture 0.75) and a single focal plane for all channels. Nuclei of the cells were segmented using CellProfiler and EGFP signal was measured over this mask. Cells were scored to be in G1-phase of the cell cycle if their integrated DNA intensity fell in the visually selected cutoff of 10-22 AU seen in non-transfected cells. Mean nuclear EGFP intensity in transfected cells was split into four equal frequency bins of increasing EGFP intensities and cell-cycle plots were drawn from these populations.

### Effect of cell-cycle on E1A protein expression in HeLa-FUCCI cells

Hela-FUCCI cells were seeded on 96-well imaging plate at a density of 7000 cells / well and grown for two days. AdV-C5 was incubated with cells at moi of ∼ 11650 virus particles per cell for 60 min at 37°C in DMEM supplemented with 0.2% BSA and penicillin-streptomycin. After removal of unbound virus, incubation was continued in DMEM supplemented with 7.5% FCS, 1% nonessential amino acids and penicillin-streptomycin for 9.5 h. Cells were fixed and stained with M58 anti-E1A (ThermoFisher Scientific, MA5-13643) and secondary anti-mouse Alexa Fluor680 (ThermoFisher Scientific, A21058) antibodies and DAPI as previously described [21]. Images were acquired with Molecular Devices automated ImageXpress Micro XL widefield imaging system using 20× S Fluor objective (numerical aperture 0.75) and a single focal plane for all channels. Nuclear stain DAPI was used to segment nuclei using CellProfiler. The segmentation output was fed to CellProfiler Analyst [113], which was used to differentiate cells into early G1, G1, G1/S and S/G2/M phases of the cell cycle. HeLa-FUCCI cells express truncated forms of Kusabira orange-fused Cdt1 as a marker for early and late G1 phases, and Geminin fused to Azami green as a marker for S and G2 phases of the cell cycle [66]. A short transition phase during the change from G1 (red) to S phase (green) is identified as G1/S phase and appears yellow due to the overlap of the two fluorescent signals. Using these set of rules, CellProfiler Analyst was trained to separate the cells into these phases and the output was re-examined to manually reassign erroneously identified phases. The software was re-trained until the output appeared satisfactory [103]. The scatter plots of mean E1A nuclear intensity expressed in different cell-cycle stage cells were plotted in JMP.

### Assaying transcriptionally active nuclear vDNAs

A549 cells were seeded at a density of 40000 on alcian blue-coated coverslips in a 24-well plate format and grown for two days. EdC-labeled AdV-C5 was incubated with cells (moi ∼ 23440 virus particles per cell) at 37°C for 60 min in DMEM supplemented with 0.2% BSA and penicillin-streptomycin. After removal of unbound virus, incubation was continued in DMEM supplemented with 7.5% FCS, 1% nonessential amino acids and penicillin-streptomycin for further 13.5 or 15 h before fixation. Acetic acid (2.5%) was added to the PFA-fixative solution when nuclear transcription sites were analyzed. RNA FISH staining with E4 intron probes (these probes detect also the E4orf1/orf2 mRNAs) and vDNA detection by click-reaction were carried out as described above. RNase A-treatment (50 µg/ml in PBS) was for 30 min at RT (control cells were kept in PBS) and cells were fixed again with 3 % PFA/PBS for 15 min at RT before proceeding to the RNA FISH and click reactions. Nuclei were stained with DAPI and and cell area with Alexa Fluor647 NHS ester. Imaging was carried out with a Leica SP5 confocal laser scanning microscope using 63× magnification oil objective (numerical aperture 1.4) and zoom factor 2.5. Stacks were recorded at 1 µm intervals with sequential acquisition using between frames switching mode and 3 or 4 times line accumulation for vDNA signals. The sensitive Leica HyD hybrid detector in the normal mode was used for vDNA signals. Maximum projections of confocal stacks and a custom-programmed CellProfiler pipeline were used to determine the E4 transcript/intron signals in the cell cytoplasm (Alexa Fluor647 cell mask minus DAPI nuclear area), as well as the number of nuclear E4 puncta or total and E4-positive vDNA numbers in the nuclear area (DAPI mask). To improve nuclear vDNA segmentation, vDNA images were processed with the Fiji plugin Rolling Ball Background Subtraction with rolling ball radius set to 5 pixels prior to running the CellProfiler pipeline. Proper cell and nucleus segmentation was controlled and adjusted manually if necessary. Proper vDNA and E4 puncta segmentations were checked from saved CellProfiler segmentation images. For setting threshold values for E4-positive vDNAs, background signal levels were first determined by placing vDNA images on E4 probe channel images of noninfected cells and the highest vDNA-associated E4 intensity was taken as the cutoff value. The selected cutoff value was compared to E4 intensity values obtained from visually selected E4-negative and –positive vDNAs to ascertain that the selected cutoff values correctly distinguished between E4-negative and –positive nuclear vDNAs. The data were sorted using KNIME Analytics Platform and cells with no nuclear vDNA signal were excluded from the analysis. GraphPad Prism was used for producing the scatter plots. Representative images were produced using Fiji as described above.

### Assaying progression of incoming vDNAs to a replication phase

A549 cells were infected with AdV-C5 at the indicated multiplicities of infection as described above. For labeling newly synthesized vDNAs, EdC (2.5 µM) was included into the culture medium for the last four hours before fixation and viral replication centers were visualized with a click-reaction using Alexa Fluor488-conjugated azide. Cytosine arabinoside (AraC; Sigma-Aldrich, C3350000, final concentration 10 µg/ml), added after removal of unbound virus, was used for suppression of viral genome replication. When probing for nuclear targets of the E1A transcript probes, RNA FISH was carried out as described above, except that 2.5% (v/v) glacial acetic acid was included into the fixative to improve detection of nuclear RNA [114] and cells were treated with Affymetrix QuantiGene ViewRNA HC screening assay kit protease (1/4000 dilution, 10 min incubation at RT) after permeabilization with Triton X-100. The protease was inactivated by incubation in the kit protease stop buffer (10min at RT). RNase A treatment for nuclear E1A probe signal was carried out after inactivation of protease as described above. S1 nuclease (ThermoFisher Scientific, EN0321) treatment was carried out after inactivation of protease with 0.56 u/µl nuclease in S1 nuclease buffer (ThermoFisher Scientific) for 60 min at 37°C, followed by two washes with PBS, incubation with 3% paraformaldehyde in PBS for 20 min at RT and two washes with PBS before proceeding to the RNA FISH staining. Control cells were incubated in S1 nuclease buffer without the nuclease. To identify incoming vDNAs that had proceeded to a replication phase, A549 cells were infected with EdC-labeled AdV-C5 (moi ∼ 23440 virus particles per cell) as described above and, after removal of unbound virus, incubation was continued at 37°C for additional 27 h with AraC in the culture medium during the last 20 h. The cells were fixed by 3% paraformaldehyde/PBS containing 2.5% acetic acid, permeabilized by Triton X-100/PBS, treated with protease and stained with E1A probes as described above, followed by incubation in ImageiT FX Signal Enhancer and click-reaction with Alexa Fluor488-conjugated azide as described above. Cells were also stained with DAPI and Alexa Fluor647 NHS ester. Imaging was carried out with a Leica SP5 confocal laser scanning microscope as described above. Representative images shown are maximum projections of confocal stacks, processed in Fiji [111] applying the same changes in brightness and contrast to all image groups in the series. Colocalization of nuclear E1A probe signal with vDNA signal was used for identification of incoming vDNAs that had proceeded to a replication phase. Alternatively, replication phase-vDNAs were identified by colocalization of anti-DBP signal with vDNA click-reaction signal. Immunofluorescence staining with mouse anti-DBP (clone A1-6; kindly provided by Nancy Reich) [115], and Alexa Fluor594-conjugated anti-mouse antibodies was carried out first, followed by a click-reaction with Alexa Fluor488-conjugated azide. Imaging was carried out with a Leica SP5 confocal laser scanning microscope as described above. Representative images shown are maximum projections of confocal stacks, processed in Fiji as described above.

### Statistical analyses

Statistical analyses were performed either by Kolmogorov-Smirnov tests using GraphPad Prism or by permutation tests using a custom-programmed R-script. Alpha factor 0.001 and 5000 permutations were used in the permutation tests. This commonly resulted in a p-value of 0. However, permutation p-values should never be zero [116]. Therefore, the p-value was calculated using the recommended formula p=(b+1)/(m+1), in which b is the number of permutations giving a difference greater than the observed difference between samples and m is the number of permutations.

## Data availability

We are going to deposit raw max projection data of images used for the scatter plot analyses, as well as CellProfiler, KNIME and R scripts to Mendeley Data once the manuscript is accepted, and we know the final contents of figures.

