## Supplementary Figures for "Cell-to-cell and genome-to-genome variability of Adenovirus transcription tuned by the cell cycle"

### S1 Figure (related to Fig. 1D): Representative images showing E1A and E1B-55K mRNAs in infected cells at 5, 8 and 12 h time points.

Images are maximum projections of confocal stacks. Scale bar = 10 µm.

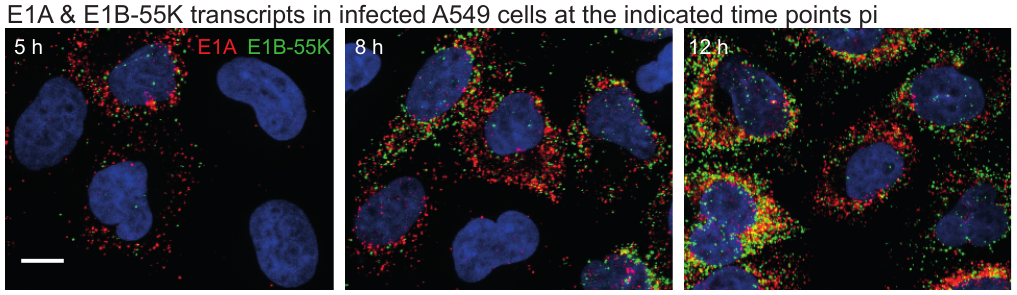

### S2 Figure (related to Fig. 2): Detection efficiency of vDNA in infected cells, poor correlation of E1A transcripts per cell with cell or nuclear areas in infected A549 cells and poor correlation of E1A transcript counts to nuclear vDNA numbers in infected HDF-TERT cells.

A) The majority of EdC-labeled AdV-C5 particles carry a detectable vDNA. EdC-labeled AdV-C5 was added to HeLa-ATCC MIB1 knockout cells at 37°C for 60 min, and, after removal of unbound virus, incubation was continued at 37°C for another 60 min before fixation. Fixed cells were stained with 9C12 anti-hexon and secondary Alexa Fluor594-conjugated anti-mouse antibodies to mark virus particles, the viral vDNA was detected by a click-reaction using azide-Alexa Fluor488 and nuclei were stained with DAPI. Images shown are maximum projections of confocal stacks. Particles carrying detectable vDNA are represented by yellow dots in the overlay image. Cell and nuclear outlines are shown. Scale bar = 10 µm. The graph shows per cell quantification of virus particles with detectable vDNA signal, one dot representing one cell. The horizontal line represents the median value. The number of cells and virus particles analyzed is indicated.

B) E1A mRNA counts were correlated to cytoplasmic and nuclear areas at single-cell level, one dot representing one cell. r_s_ denotes the Spearman’s correlation rank coefficient. No significant correlation was was observed between nuclear area and E1A transcript counts. C) Comparison of E1A mRNA abundancies and nuclear vDNA counts at single-cell level in HDF-TERT cells infected with EdC-labeled AdV-C5. The virus was incubated with cells at 37°C for 15 h, and, after removal of unbound virus, incubation was continued at 37°C for additional 7 h. Fixed cells were analyzed as described in legend to Fig. 2. Number of cells analyzed was 29. No significant monotonic correlation between the nuclear vDNA and E1A transcript counts was observed (Spearman’s correlation test).

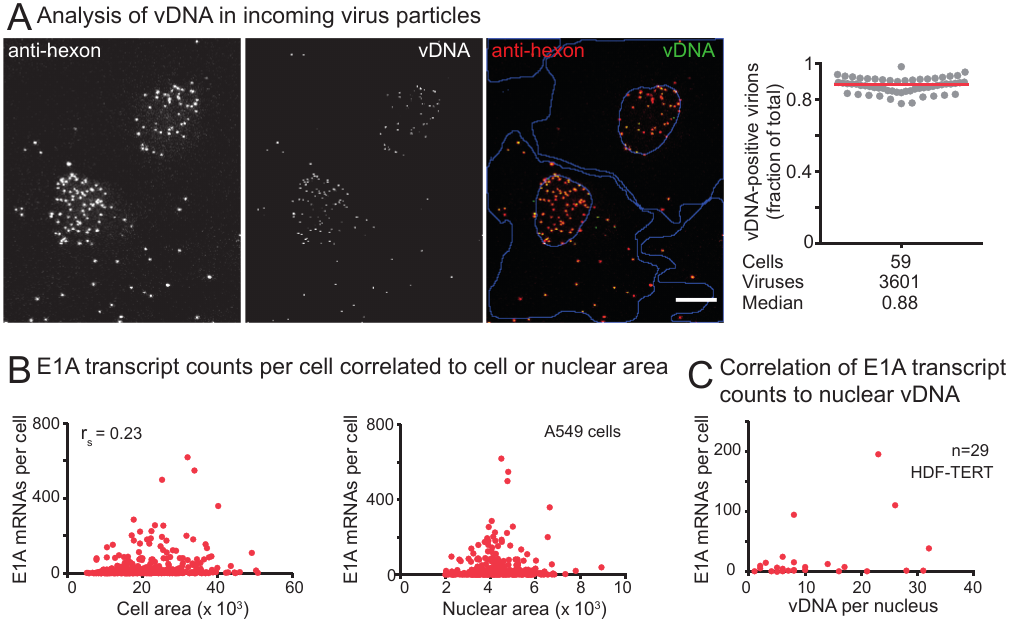

### S3 Figure (related to Fig. 3). Effect of host cell-cycle stage on accumulation of E1A transcripts early in infection.

A) Infection in HeLa-FUCCI suggests that E1A mRNAs accumulate more rapidly in G1 cells than in S/G2/M cells. AdV-C5 was added to HeLa-FUCCI cells (moi ~ 11650 virus particles per cell) at 37°C for 60 min, and, after removal of unbound virus, incubation was continued at 37°C for additional 6.5 h before fixation. Fixed cells were stained with E1A bDNA-FISH probes. In general, G1 cells (Cdt1-positive, Geminin-negative) had accumulated more E1A transcripts than Geminin-positive cells (S/G2/M cells), but the spectral overlap of E1A and Cdt1 signals precluded quantitative assessment of the experiment. Images shown are maximum projections of confocal stacks. Scale bar = 10 µm.

B) Cell-cycle histograms of noninfected and infected A549 cells at 4 h post infection drawn from integrated nuclear DAPI intensities of the cells. The visually selected cutoff values for G1 cells (132 – 200 arbitrary units) is indicated as a shaded area. Similarities between the noninfected and infected cell histograms indicate that at this time point post infection the virus has not yet induced large scale progression of the host cells to the S-phase.

C) Cells with high amounts of E1A transcripts at 7 h post infection are predominantly G1 cells. Infection was carried out as described in legend to Fig.3B, except that cells were analyzed at 6 h after removal of unbound virus. The mean cytoplasmic E1A probe signal intensities were used for estimation of E1A transcript abundancies per cell and the results are shown as a boxplot (left-hand panel). Outliers for E1A expression, i.e. cells with mean cytoplasmic E1A intensities more than 1.5 × interquartile range from the 75^th^ percentile, are colored black, and, as shown in the right-hand cell-cycle histogram, these outlier cells are predominantly in the G1 cell-cycle phase. Overall, about 57.2 % of cells in the total population were G1 cells, whereas the percentage of G1 cells in the outlier population was 72.6 %.

D) Cell-cycle histograms of non-infected or infected A375 cells continuously incubated in serum-free medium (starved + starved) or of cells that were switched to serum-containing medium after removal of unbound AdV-C5 (starved + serum). The visually selected cutoff values for G1 cells is indicated as a shaded area and the percentage of G1 cells in the different cell populations is shown.

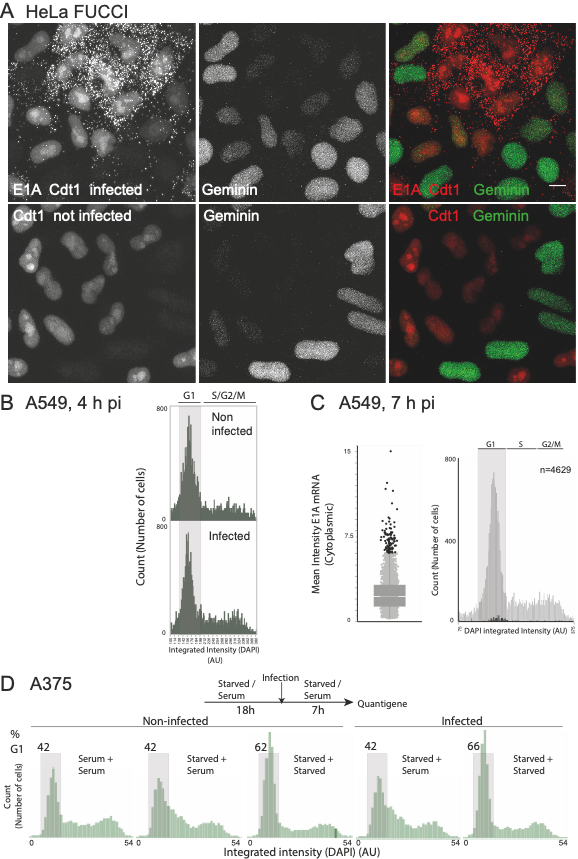

S4 Figure (related to Fig. 5): Cell-cycle histogram of cells included in the Fig. 5 data.

The histogram was drawn from integrated nuclear DAPI intensities of 523 cells and the visually selected cutoff (1100 – 1800) for G1 cells is indicated.

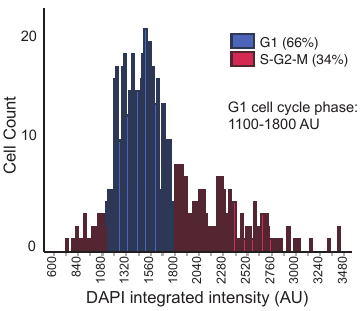

### S5 Figure (related to Fig. 6): E4 orf1/orf2 abundance and correlation with incoming vDNA genomes.

A) Cytoplasmic transcript counts for E4orf1/orf2 mRNAs. A549 cells were infected with EdC-labeled AdV-C5 as described in legend to Fig. 2 except that cells were fixed and stained with bDNA-FISH probes targeting the E4 transcription unit at 15 h or 17 h post removal of unbound virus. The probes recognize E4-derived intron sequences as well as mature E4orf1 and E4orf2 mRNAs. The target sequence on E4orf2 mRNAs is shorter than on E4orf1 transcripts, but the signal from E4orf2 transcripts is most likely visible as well. The scatterplot shows quantification of cytoplasmic E4orf1/orf2 transcripts, one dot representing one cell. NI indicates noninfected control cells. Horizontal bars represent median values and the number of cells analyzed is indicated.

B) Nuclear E4 probe dots at 16h post infection mostly originate from RNA. A549 cells were infected with EdC-labeled AdV-C5 as described in legend to Fig.2 and 15 h post removal of unbound virus cells were fixed and stained with the E4 bDNA-FISH probes. Acetic acid was included into the fixative solution to improve accessibility of probes to nuclear targets, but acetic acid suppresses signals from cytoplasmic targets and therefore cells appear to be devoid of cytoplasmic E4 probe signals. RNase A denotes samples treated with RNase A prior to the FISH staining. The images shown are maximum projections of confocal stacks. Nuclei outlines were drawn from DAPI-stained nuclei. Scale bars = 10 µm.

C) Quantification of nuclear E4 mRNA puncta at 16 h post infection in noninfected and infected cells, and infected cells treated with RNase A. CellProfiler was used to score the number of nuclear E4 dots from maximum projection images (one dot represents one nucleus). Horizontal bars represent median values and the number of cells analyzed is indicated. Cells with RNase A-resistent puncta most likely represent cells in which vDNAs have already progressed into a replication phase (see also S6 Fig).

D) vDNAs within the same nucleus display heterogeneous transcriptional activity. A549 cells were infected with EdC-labeled AdV-C5 and processed as described in legends to Fig. 2 and Fig. 6, except that cells were fixed 13.5 h post removal of unbound virus. vDNAs localizing over DAPI-mask in maximum projection images were counted as nuclear vDNAs and colocalization of these vDNAs with E4 bDNA-FISH probe signals was taken to indicate transcriptionally active vDNAs. The scatterplot correlates total nuclear vDNA numbers to the number of E4 signal-positive vDNAs within the same nucleus, one dot representing one nucleus. Number of cells analyzed was 140.

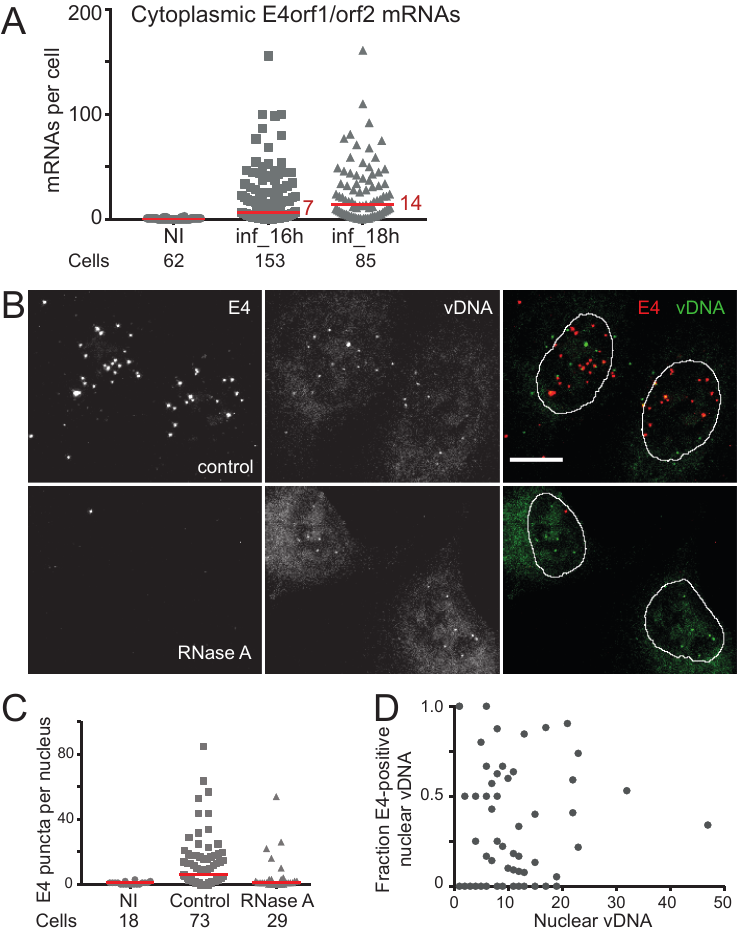

### S6 Figure (related to Fig. 6): E1A bDNA-FISH probes stain viral replication centers late in infection.

A) AdV-C5 was added to A549 cells at 37°C for 60 min (moi ~ 46600 virus particles per cell), and, after removal of unbound virus, cells were further incubated for 17 h with EdC present in the incubation medium during the last 4 h. Fixed cells were stained with E1A bDNA-FISH probes and the newly synthesized EdC-labeled vDNA in the viral replication centers was detected by a click-reaction using azide-Alexa Fluor488. The EdC-signal in infected cells displayed the characteristic flower-like pattern of viral replication centers, whereas a smoother nuclear staining pattern was observed for noninfected cells. In the AraC-treated sample, AraC was added to the culture medium after removal of unbound virus and incubation was continued for 17 h. The E1A probes yield punctate pattern in AraC-treated cells, the puncta most likely representing stalled viral replication forks [77].

B) The E1A bDNA-FISH probes hybridize to nuclear single-stranded vDNA late in infection. In the experiment shown in the upper panel, AdV-C5 was added to A549 cells at 37°C for 60 min (moi ~ 40800 virus particles per cell), and, after removal of unbound virus, incubation was continued at 37°C for additional 11 h in the presence of AraC. Fixed cells were treated or not with RNase A prior to staining with E1A bDNA-FISH probes. Nuclear E1A puncta were resistant to the RNase A-treatment, whereas similar amounts of RNase A efficiently degraded cytoplasmic E1A mRNAs (Fig. 1B). In the experiment shown in the lower panel, AdV-C5 was added to A549 cells (moi ~ 27200 virus particles per cell) at 37°C for 60 min, and, after removal of unbound virus, incubation was continued at 37°C for additional 27.5 h before fixation, with AraC present in the medium during the last 23.5 h. Fixed cells were treated or not with S1 nuclease prior to staining with E1A bDNA-FISH probes. S1 nuclease degrades single-stranded nucleic acids and the nuclear E1A signals were suppressed by this nuclease treatment. Sensitivity of nuclear E1A signal to S1 nuclease, but not to RNase A, indicates that the nuclear E1A probe signals originate from single-stranded vDNA. Cytoplasmic E1A mRNAs are not well visible in the images because acetic acid was included into the fixative solution to improve accessibility of probes to nuclear targets but this fixation condition suppresses signals from cytoplasmic targets. All images shown are maximum projections of confocal stacks. Nuclei (pseudocolored blue) were stained with DAPI. Scale bars = 10 µm.

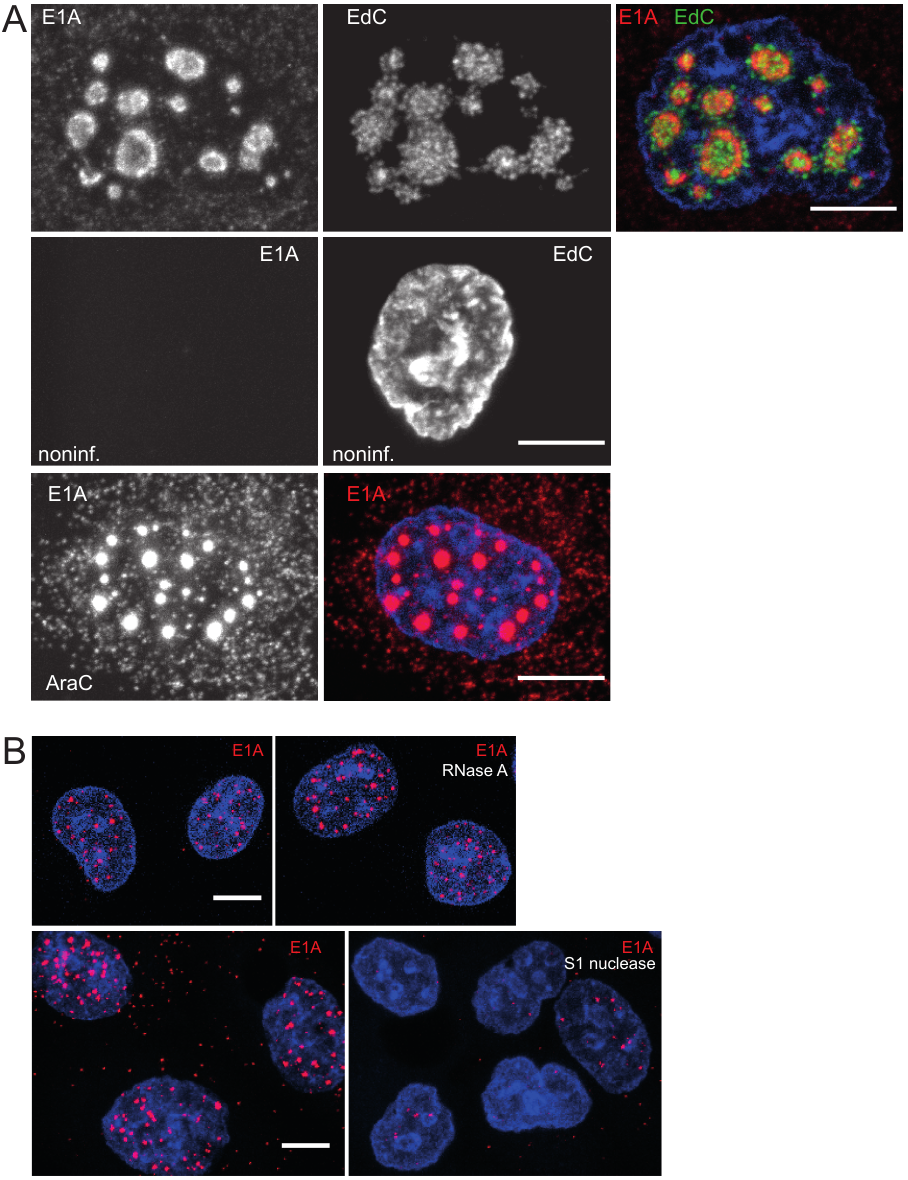
