## Supplementary material for "Cell-to-cell and genome-to-genome variability of Adenovirus transcription tuned by the cell cycle": Key resources

**KEY RESOURCES TABLE**

| REAGENT or RESOURCE | SOURCE | IDENTIFIER |
| --- | --- | --- |
| **Antibodies** | | |
| Anti-hexon (9C12) | L. Fayadat-Dilman/W. Olijve  University of Iowa, Developmental Studies Hybridoma Bank | TC31-9C12.C9 |
| Anti-E1A (M58) | ThermoFisher Scientific | Cat #MA5-13643 |
| Anti-DBP | Clone A1-6; Kindly provided by Nancy Reich | N/A |
| Alexa Fluor488-conjugated goat anti-mouse | ThermoFisher Scientific | Cat #A11029 |
| Alexa Fluor594-conjugated anti-mouse | ThermoFisher Scientific | Cat #A21203 |
| Alexa Fluor680-conjugated anti-mouse | ThermoFisher Scientific | Cat #A21058 |
| **Virus Strains** | | |
| AdV-C5 | (V. Prasad et al., 2014) | N/A |
| **Chemicals, Peptides, and Recombinant Proteins** | | |
| Alexa Fluor647 NHS-Ester | ThermoFisher Scientific | Cat #A20006 |
| Alexa Fluor647-conjugated wheat germ agglutinin | ThermoFisher Scientific | Cat #W32466 |
| Alexa Fluor488-conjugated azide | ThermoFisher Scientific | Cat #A10266 |
| Quantigene ViewRNA high content screening assay kit | ThermoFisher Scientific | QVP0011, QVP0201/LP1-550,  QVP0201/LP4-488,  custom-made probes |
| S1 nuclease | ThermoFisher Scientific | Cat #EN0321 |
| ImageiT FX Signal Enhancer | ThermoFisher Scientific | Cat #I36933 |
| 5-ethynyl-2'-deoxycytidine (EdC) | Sigma-Aldrich | Cat #T511307 |
| Bovine Serum Albumin (BSA) | Sigma-Aldrich | Cat #A9418 |
| Lipofectamine 2000 | Invitrogen | Cat #11668019 |
| Optimem low-serum medium | ThermoFisher Scientific | Cat #11058021 |
| Penicillin-streptomycin | Sigma-Aldrich | Cat #P0781 |
| DNeasy Blood and Tissue kit | Qiagen | Cat #69506 |
| Cytosine arabinoside (AraC) | Sigma-Aldrich | Cat #C3350000 |
| **Experimental Models: Cell Lines** | | |
| Human: HeLa ATCC | American Type Cell Culture | [Cat](http://www.lgcstandards-atcc.org/products/all/CCL-2.aspx?geo_country=ch) #CCL-2 |
| Human: HeLa ATCC MIB1 knockout cells | Michael Bauer | <https://doi.org/10.1016/j.celrep.2019.11.064> |
| Human: HeLa FUCCI | Atsushi Miyawaki | <https://doi.org/10.1016/j.cell.2007.12.033> |
| Human: A375 | American Type Cell Culture | Cat #CRL-1619 |
| Human: A549 ATCC | American Type Cell Culture | Cat #CCL-185 |
| Human: A549 | Our lab’s old A549 clone |  |
| Human: HDF-TERT | Patrick Hearing,  Kathleen Rundell | <https://doi:10.1006/viro.2001.1204> |
| **Recombinant DNA** | | |
| pN1-E1A-EGFP | Sten Strunze/Greber lab | N/A |
| pN1-CMV-EGFP | V. Prasad et al., Nature Communications 2020, in press |  |
| **Software and Algorithms** | | |
| Cell Profiler | (Carpenter et al., 2006) | Version 3 |
| KNIME | KNIME Analytics Platform | https://www.knime.org/knime-analytics- platform |
| JMP | SAS | Version 13 |
| Graphpad Prism | GraphPad Software, Inc. La Jolla | Version 8 |

Table 1. List of primers used in the study

| **Primers** | | |
| --- | --- | --- |
| E1A-forward (ChIP) | V. Prasad et al., Nature Communications, 2020, in press | GGTGGAGTTTGTGACGTGG |
| E1A-forward (ChIP) | V. Prasad et al., Nature Communications, 2020, in press | CGCGCGAAAATTGTCACTTC |
